# Linguistic and attentional factors – not statistical regularities – contribute to word-selective neural responses with FPVS-oddball paradigms

**DOI:** 10.1101/2023.02.20.528973

**Authors:** Aliette Lochy, Bruno Rossion, Matthew Lambon-Ralph, Angélique Volfart, Olaf Hauk, Christine Schiltz

## Abstract

In recent years, a fast periodic oddball-like paradigm has proved to be highly sensitive to measure category-selective visual word representation and characterize its development and neural basis. In this approach, deviant words are inserted in rapid streams of base stimuli every n^th^ occurrence (e.g., Lochy et al., 2015). To understand the nature of word-selective representation and improve its measurement, we tested 22 adults with EEG, assessing the impact of discrimination coarseness (deviant words among nonwords *or* pseudowords), the relative frequency of item repetition (set size *or* item repetition controlled for deviant vs. base stimuli), and the nature of the orthogonal attentional task (focused or deployed spatial attention). In all stimulation sequences, base stimuli were presented at 10 Hz, with words inserted every 5 stimuli generating word-selective responses in the EEG spectra at 2 Hz and harmonics. Word-selective occipito-temporal responses were robust at the individual level, left-lateralized and sensitive to wordlikeness of base stimuli, being stronger in the coarser categorical contrast (among nonwords). Amplitudes were not affected by item repetition, showing that implicit statistical learning about a relative token frequency difference for deviant stimuli does not contribute to the word-selective neural activity, at least with relatively large stimulus set sizes (n=30). Finally, the broad attentional deployment task produced stronger responses than a focused task, an important finding for future studies in the field. Taken together, these results confirm the linguistic nature of word-selective responses, strengthen the validity and increase the sensitivity of the FPVS-EEG oddball paradigm to measure visual word recognition.

**Highlights:**

- Word-selective responses measured in fast periodic visual stimulation with EEG are linguistic in nature
- Word-selective responses reflect prelexical or lexical processes depending on the contrast (words in nonwords or pseudowords respectively)
- Using sufficiently large sets (30 items) prevents the extraction of statistical regularities and hence, statistical learning
- Using an orthogonal task involving broad, rather than focused, spatial attention increases amplitude of the neural responses
- Sensitivity of the paradigm to detect significant responses at the individual level is very good (95% for prelexical and about 80% for lexical word responses)

## Introduction

In recent years, a frequency-tagging electroencephalographic (EEG) approach based on ‘oddball’, or ‘oddball-like’ design, has been successfully applied to explore visual recognition in the nonverbal (objects, faces, or numerosities; e.g., Guillaume et al., 2018, 2020; Liu-Shuang et al., 2014a; Marinova et al., 2021; Marlair et al., 2021; Nurdal et al., 2021; Retter et al., 2020; Stothart et al., 2017) and verbal domains (letters, words, word semantic categories; e.g., (Lochy et al., 2015; Volfart et al., 2021). This fast periodic visual stimulation (FPVS) oddball approach is typically based on variable exemplars of a frequent stimulus category (or “base” category) presented at a rapid rate (e.g., 10 Hz), interrupted at a slower periodic rate (usually 1/5, e.g., 2 Hz) by a contrastive stimulus category (or “deviant” category) (Rossion et al., 2018, 2020 for reviews). If the deviant category (e.g., words) is discriminated from the base category (e.g., non-words or pseudo-words, NW and PW hereafter), it elicits a neural response measurable in the frequency domain at exactly the frequency of the deviant category (e.g., 2Hz) (Lochy et al., 2015). This approach has substantial advantages in terms of objectivity as responses occur at experimentally predefined frequencies. Furthermore, it is characterized by great sensitivity: because of its high signal-to-noise ratio (SNR), the neural response can be measured in various sensitive populations such as infants, children or patients (e.g., de Heering & Rossion, 2015; Lochy et al., 2016, 2019; Peykarjou et al., 2017; Vettori et al., 2019) in only a few minutes of recordings. Finally, neural responses are measured without requiring any explicit task, another advantage especially when collecting data in developmental or clinical populations.

Following the first study in French (Lochy et al., 2015), the FPVS-EEG oddball approach was used to measure category-selectivity to visual letters and words. It has revealed left hemispheric cortical specialization for letters in young pre-readers related to their letter knowledge (Lochy et al., 2016), neural changes occurring over one year in beginning readers (van de Walle de Ghelcke et al., 2021), impact of teaching methods for learning to read (van de Walle de Ghelcke et al., 2020) and the neural basis of letter-selective and (pre)lexical responses in the left ventral occipito-temporal cortex with intracerebral recordings (Lochy et al., 2018). It has also recently been successfully extended to German (Aristei et al., 2017), though evidence of its suitability to English is currently lacking (Barnes et al., 2021; Wang et al., 2021)

Given FPVS-EEG’s wide applicability, the main objective of the current study is to understand the nature of word-selective neural responses elicited in this approach. In particular, it is crucial to ascertain that the categorical responses measured with this paradigm stem from a higher conceptual (linguistic) level, and not from potential confounding peripheral processes or design factors. Specifically, we explore how discrimination coarseness and item repetition rate (implying potential statistical learning) modulate responses to words. A second aim was also to explore if the type of orthogonal attention task, spatially focused or deployed, would influence amplitude of responses. Finally, a last objective was to evaluate if this approach is sufficiently sensitive at the individual participant level. Below, we review these points in turn.

To understand the nature of the neural response to words in FPVS, we first investigate the impact of discrimination coarseness between the two contrasted categories. It is important to bear in mind that the response to the deviant stimuli in this paradigm is not measured in absolute terms, but as an index of a *differential* processing between base and deviant stimuli (i.e., all common processes between base and deviant stimuli project to the common base rate response, which can be also objectively identified in the EEG spectrum). Thus, we do not measure an absolute response to the category of words, but a response to words among another type of stimuli, and hence, the nature of the process at hand is *constrained* by the type of stimuli used. In the domain of written word recognition, the deviant response is modulated in amplitude and scalp topography by the (dis)similarity between the two categories that are contrasted (Lochy et al., 2015). As an example, when words are discriminated among pseudofonts, the response is larger and more bilateral (although with a left hemispheric dominance) than for words among nonwords (NW) or pseudowords (PW) (Lochy et al., 2015). Indeed, discriminated against pseudo-letters, responses to written words may reflect not only lexical and orthographic processes but also, more basically, the contrast of real letters to non-existing letter-like forms, thus generating more bilateral, lower-level, visual responses. Similarly, words among NW appear to trigger a larger amplitude response than words among PW (Lochy et al., 2015), and this could be because this coarser contrast may be based on pre-lexical orthographic plausibility detection by comparison to the finer contrast. Here, we further explored the sensitivity of discrimination responses for words to the level of discrimination induced by the categorical contrast (NW or PW, Figure 1A), using a different stimulus set than the princeps study (Lochy et al., 2015) and a larger sample of participants (N=10 vs. 22).

**Figure 1.**
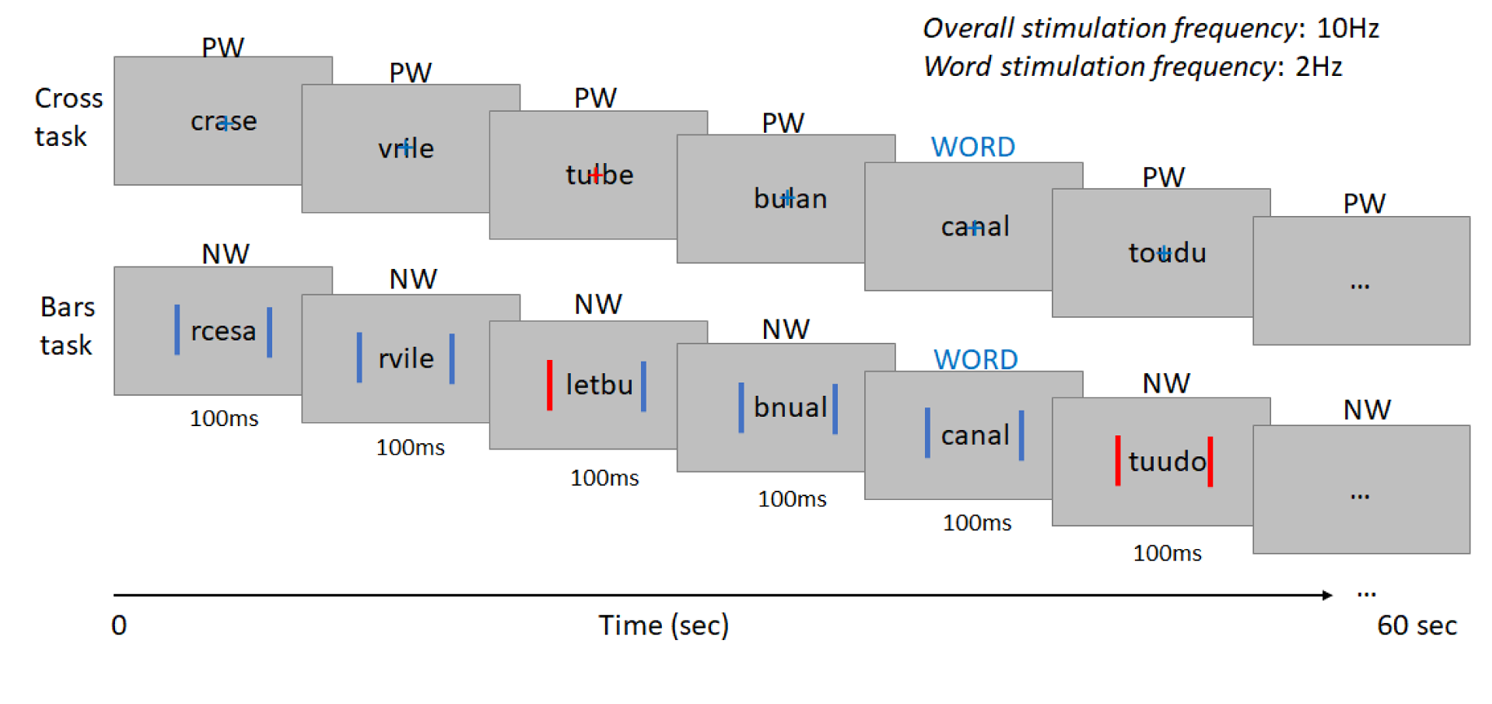
Experimental design. Sequences of letter-strings were presented at 10Hz (10 stimuli/sec) during 60 seconds. Deviant stimuli (words) were inserted every five items (at 2Hz) among two types of base stimuli: pseudowords (PWW, top row) or nonwords (NWW, bottom row), inducing a fine-grained or a coarse discrimination contrast. Relative exemplar frequency of base and deviant stimuli was manipulated (*not represented in the Figure*), controlling either item repetition (with different set sizes for base and deviant stimuli), or set size (with different number of item repetitions for base and deviant stimuli). Two different orthogonal attentional tasks were implemented, requiring to monitor colour-changes either to a central cross (top row), or vertical bars (bottom row) flanked left and right of the letter-strings. Participants had to respond when the cross turned red, or when both bars turned red at the same time. Each type of attentional task (bars/cross) was performed on each discrimination type (NWW/PWW) and each control type (set size/exemplar repetition), resulting in a 2 x 2 x 2 design.

Responses to deviant words in this paradigm might however stem not (only) from linguistic processes (words vs nonwords/pseudowords) but from relative item repetition differences between base and deviant categories. Due to the nature of the oddball paradigm, there is usually an inherent difference between the rare and frequent sets of stimuli in the number of exemplar repetitions, or *relative item frequency*. Indeed, when two sets of an identical number of stimuli are used (e.g., 30 per set) and if the deviants are presented every five items (1/5), then the relative frequency of each *item* differs across categories (e.g., Lochy et al., 2015). Over a certain period, the base stimuli are presented 4 times more often than the deviants. For a concrete example, if the base stimulation frequency is at 10Hz (10 images per second) with a deviant every 5 items as in Lochy et al. (2015), over 1 minute of stimulation, a total of 600 images are presented among which are 480 base stimuli and 120 deviants. If the set sizes are equivalent, e.g., 30 items per set, then each base stimulus is repeated 16 times while each deviant stimulus is repeated only 4 times. Thus, with this paradigm, either the number of exemplars per set is matched (and item repetition is not, as in most previous studies) or the number of exemplar *repetitions* is matched (and the number of stimuli per set differs).

Given this, an important question is whether the relative item frequency can contribute, at least in part, to the measured neural discrimination response. That is, one cannot exclude that items of the deviant set would be discriminated, not because of their experimentally manipulated properties (e.g., the psycholinguistic contrast between words and non-words), but merely because they are repeated less often in the course of stimulation sequences (and the whole experiment). Indeed, since the seminal studies in language acquisition that revealed fast learning of the statistical structure of meaningless syllable sequences varying in transitional probabilities in both adults and infants (Saffran, Aslin, et al., 1996; Saffran, Newport, et al., 1996), the sensitivity of our cognitive and neural system to detect structured patterns of the environment, also called *implicit statistical learning*, has been widely documented in a variety of domains (e.g., Frost et al., 2019). This powerful mechanism has been suggested to be at the root of word segmentation (Saffran, Aslin, et al., 1996; Saffran, Newport, et al., 1996), vocabulary acquisition (Shafto et al., 2012), complex sentences processing and grammar learning (Dienes et al., 1991; Misyak et al., 2010; Misyak & Christiansen, 2012), syntactic comprehension (Kidd & Arciuli, 2016) and reading acquisition (Arciuli & Simpson, 2012). Therefore, in FPVS-EEG studies, one cannot exclude the possibility that our brain can extract the statistical regularities in the stimuli distribution and relative item frequency, and the patterned periodicity of occurrence of deviants among base stimuli, generating or modulating oddball responses to deviant stimuli independently of their categorical nature. Intriguingly, two recent FPVS-EEG studies actually suggested that it could be the case, showing that arbitrary groupings of digits (Guillaume et al., 2020) or of letter-strings (de Rosa et al., 2022) gave rise to some discrimination response, presumably due to the difference in relative item frequency, as there was no a priori reason for them to be discriminated from each other.

Here, to investigate the potential impact of the relative exemplar frequency within base/deviant categories on neural responses to deviant words, we contrasted stimulation sequences in which set size was controlled, and the number of exemplar repetitions differed between base and deviant (i.e., 30 deviants-repeated 4 times each in a sequence, and 30 base stimuli repeated 16 times each in a sequence), to stimulation sequences in which the number of item repetitions, but not the set size, was controlled (i.e., 30 deviant and 120 base stimuli, all repeated 4 times in a sequence). Several hypotheses can be formulated. First, if implicit statistical learning about exemplar repetition is the only source of the neural oddball response, then discrimination of deviants should occur *only* when the set size is controlled (and item repetition rate is not). Such an observation would be in line with the significant discrimination occurring between sets that do not differ linguistically (de Rosa et al., 2022), and would imply that the linguistic status of words is not a key variable in FPVS-EEG responses, thus that these responses might have been wrongly interpreted in previous studies (Lochy et al., 2015, 2018; Volfart et al., 2021). *No response* should occur when item repetition rate is controlled, as there would be no difference in relative item frequency between words and base stimuli, as was observed for English words among pseudowords (Barnes et al., 2021). Alternatively, the neural oddball response to words may have *nothing* to do with statistical learning of regularities, so that discrimination of deviants would occur equally strongly in the two design types. Finally, *both* statistical learning and high-level word-selectivity could contribute to the neural discrimination. In that case, a quantitative difference between the two conditions should be observed, with discrimination responses being larger when set size is controlled (and item repetition differs), but still present when item repetition is controlled (and set size differs).

The third objective of this study was to assess whether there is an influence of the orthogonal attentional task performed while participants watch the stream of stimuli on the screen during FPVS. In most previous FPVS-EEG studies with complex stimuli, attention is directed towards changes of colour to a fixation cross in the centre of the screen (since Rossion & Boremanse, 2011). This means that discrimination responses to deviant stimuli do not depend on engagement of cognitive resources on the stimuli per se. However, the spatial distribution of this attention may well allow better/worse processing of the stimuli. As used with letter strings, a central cross task might have primarily focused attention on the central/foveal letters (Nazir et al., 2004; Yeshurun & Carrasco, 1998) not located at the optimal fixation point for visual word recognition (*preferred landing position*, Rayner, 1998). Moreover, a fixation cross superimposed on letters partly masks 1 or 2 letters, depending on stimulus size (Figure 1). Taking into account this issue, here we assess the impact of two different tasks requiring either focused or deployed spatial attention. The latter was implemented by colour-monitoring of two simultaneous vertical bars left and right from the letter string (see Figure 1; e.g., as used recently in Yan et al., (2022) with face stimuli). We expect that this alternative task might induce processing of a larger portion of the visual field and thus increase attention to all letters presented, as suggested by studies showing that the size of attentional focus is adjusted to the size of the spatial cue (Eriksen & St James, 1986; Turatto et al., 2000) and may improve perceptual identification (Posner, 1988). Furthermore, some findings suggest that the impact of broadening attention by cueing both sides of the target might have a differential impact depending on lexicality of targets, being more beneficial for frequent words than pseudowords (Montani et al., 2014); although see (Ducrot & Grainger, 2007), who suggest no effect of spatial cueing on centrally presented targets). If this is the case even when no explicit identification is required, we expect the deployed attention task to give rise to larger word-selective responses than the focused task. Furthermore, this increase of responses should be selective to discrimination (because of enhanced processing) and should not influence general visual responses at the base stimulation rate.

Our final objective was to evaluate the sensitivity of the paradigm to elicit responses at the individual participant-level. Indeed, in the princeps study of (Lochy et al., 2015), 8 participants out of 10 showed a significant response to words among nonwords or pseudowords. This was recently challenged in another, less orthographically-transparent language (English) where only 10% of datasets were significant (10 participants, Barnes et al., 2021). Therefore, we examined this issue again on a larger sample of participants and different set of stimuli, as well as separately for each discrimination level (words in nonwords/ pseudowords).

## Material and Methods

### Participants

Twenty-four adults with normal/corrected-to-normal vision were tested after they gave their written informed consent for a study approved by the Ethical Committee of the University of Luxembourg. Participants were neither aware of the goal of the experiment, nor that a change of stimulus type occurred at a periodic rate during stimulation. Two participants were excluded because their EEG data was too noisy and thus analyses were performed on 22 participants (11 males, mean age = 23.54 years; range = 20-29 years).

### EEG testing stimuli

Thirty French words (concrete frequent nouns) were chosen (Supp.Mat Table S1). All were 5 letters long and presented in their singular form. Lexical frequency (Celex database in WordGen (Duyck et al., 2004) was on average 86.77 per million (min: 13.42, max: 734.51, st.dev.: 129.90). 17 words were monosyllabic, 12 bi-syllabic and 1 contained three syllables. For the fine-grained lexical discrimination level, four different sets of 30 pseudowords were created to match the words pairwise in consonant-vowel structure, number of syllables, number of orthographic neighbours and bigram frequency. Each set of PW did not differ from words on these variables (see Table 1, all stimuli in Supp Mat). For the coarse-grained pre-lexical discrimination level, four different sets of nonwords were also created. To do so, letters constituting each PW were shuffled in order to give rise to a (mostly) non-pronounceable item, matched in letter identity only with PW and purposely not matched with words in number of orthographic neighbours or bigram frequency. Characteristics of these 4 sets of 30 non-words are described in Table 1. In total, there were thus 30 words, 120 PW and 120 NW.

**Table 1.**
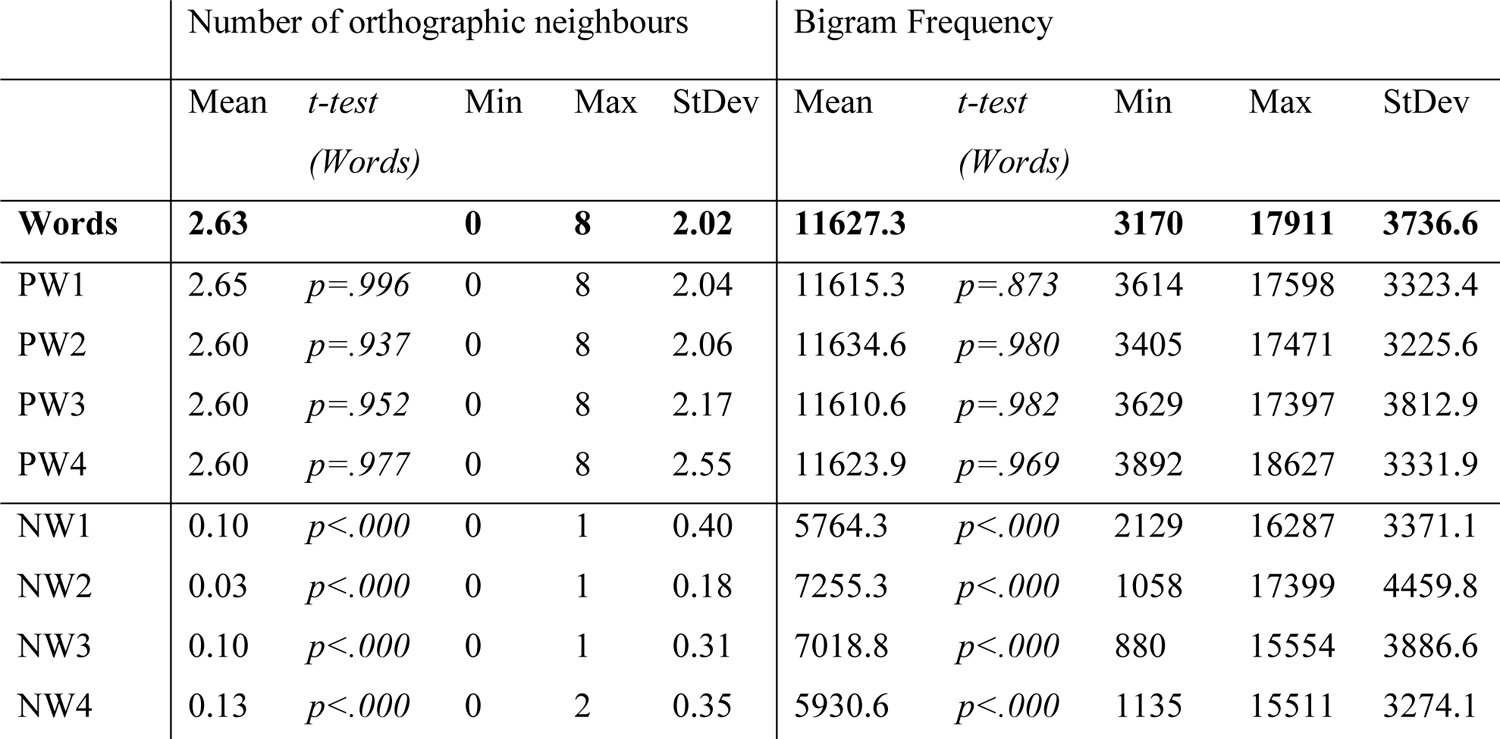
Stimulus characteristics: mean, min, max and standard deviation for the number of orthographic neighbours (first column) and for bigram frequency (second column) for words and each set of pseudowords (PW) and nonwords (NW). The p-values of t-tests comparing each list of PW and NW to words show that PW did not differ, while NW differed on these two parameters from words.

Final stimuli were maximum 238 x 71 pixels. At a distance of 1 m, displayed with a 1920 x 1080 pixel resolution, they had an average size of 3.97 x 1.26 degrees of visual angle. Stimuli were presented at the centre of the screen with no immediate repetition of the same stimulus.

### Procedure

Each stimulation sequence started with the fixation corresponding to the task (cross or vertical bars) for 2 – 5 seconds, 2 seconds of gradual stimulation fade in, 60 seconds of the stimulation sequence, and 2 seconds of gradual fade out (see Fig. 1). As in previous studies (e.g., Lochy et al., 2015), stimuli were presented by means of sinusoidal contrast modulation at a base frequency rate of 10Hz (i.e., one item every 100 ms) (Figure 1). Stimuli were presented with Java SE Version 8. Every sequence had the same structure: stimuli of the base category were presented at 10Hz (non-words or pseudo-words), and every fifth item was a deviant stimulus (words, frequency of 2Hz, thus every 500 ms).

Four conditions were presented, in a 2×2 design crossing Discrimination Level (coarse/fine) and Control Type (set size controlled/ item repetition controlled). In the *coarse discrimination level*, words were presented every five items among strings of non-words (NW-W); while in the *fine-grained discrimination level*, words were presented every five items among strings of pseudowords (PW-W). *Set size* was controlled when, as per the original study (Lochy et al., 2014), 30 words were presented as deviants among 30 base stimuli (nonwords or pseudowords). However in that case, the number of repetitions is much higher for the base stimuli: at 10Hz, 600 items are displayed in a 60-sec sequence, among which 480 base stimuli (30 PW or NW x 16) and 120 deviants (30 W x4). *Item repetition* was controlled when 30 words were presented among 120 different base stimuli (either nonwords or pseudowords). In that case, each item was presented an identical number of times (4x) during each 60-sec sequence, but the size of sets differed.

Two orthogonal tasks were designed to assess the impact of focus or deployed spatial attention (Figure 1). The focus attention required to monitor a colour change occurring randomly on a cross in the centre of the screen, while stimuli were displayed, by pressing the space bar upon each colour-change (blue-to-red). The deployed attention required to monitor simultaneously two vertical bars displayed left and right of the stimuli (−0.3 and 0.3 in unit coordinates), where colour changes could occur either on the left, on the right, or on both at the same time, and to press the space bar when both vertical bars changed colour simultaneously. In each task, there were 8 colour-changes per 60s sequence. While for the cross, all colour-changes required a key press (“go” trials), for the bars, approximately half of the trials required a key press (“go” trials; 4.05/8) and half did not (“no-go” trials; 3.95/8). These two tasks were presented by block and their order was counterbalanced across participants.

Each sequence was repeated twice in each task, for a total of 4 conditions x 3 repetitions x 60 seconds per task. A pause was taken between each of the 24 sequences, which were initiated manually to ensure low-artefact EEG signals.

### EEG acquisition and preprocessing

Participants were seated comfortably at 1 m from the computer screen in a quiet room of the University. EEG was acquired at 1024Hz using a 68-channel Biosemi Active II system (Biosemi, Amsterdam, Netherlands), with electrodes including 64 channels standard 10-20 system locations (http://www.biosemi.com) plus a row of posterior electrodes including PO9, I1, I2, PO10. The magnitude of the offset of all electrodes, referenced to the common mode sense (CMS), was held below 50 mV. EEG analyses were carried out using Letswave 6 (https://github.com/NOCIONS/letswave6), and Matlab 2012 (The Mathworks). After FFT band-pass filtering between 0.1 and 100Hz, EEG data were segmented to include 2 seconds before and after each sequence, resulting in 64-second segments (−2 – 62 s). Data files were then resampled to 512Hz to reduce file size and data processing time. Artefact-ridden or noisy channels were replaced using linear interpolation (on average, 1.54% of channels). All channels were re-referenced to the common average. EEG recordings were then segmented again from stimulation onset until 59.998 seconds, corresponding to the largest number of complete cycles of 500 ms (120 cycles) at the 2 Hz frequency within the 60 seconds of stimulation period. The data were analysed following the same procedure as in previous studies with similar paradigms (e.g., (Lochy et al., 2015, 2016) but is nevertheless detailed here.

### Frequency domain analysis

Per condition, the three stimulation sequences were averaged in the time domain for each participant, in order to increase SNR. A Fast Fourier Transform (FFT) was applied to the averaged time-window, and normalized amplitude spectra were extracted for all channels. This yielded EEG spectra with a high frequency resolution (1/59.998 s = 0.016Hz), increasing SNR and allowing unambiguous identification of the response at the exact frequencies of interest (i.e., 10Hz for the base stimulation rate and 2Hz and harmonics for the other category stimulation). Given that our response of interest falls into a strictly defined frequency bin (related to stimulation frequency), we considered surrounding bins as noise, or baseline. The latter is defined as the 20 surrounding bins of each target bin, excluding the immediately adjacent and the extreme (min and max) bins (Liu-Shuang et al., 2014b; Rossion et al., 2012; Srinivasan et al., 1999). We then computed three indices. First, the signal-to-noise ratio (SNR) was estimated across the EEG spectrum, by dividing the amplitude at each frequency bin by the average amplitude of 20 surrounding bins (10 on each side; Liu-Shuang et al., 2014b). Second, to quantify the responses of interest in microvolts, the average voltage amplitude of the 20 surrounding bins (i.e., the noise) was subtracted out (Retter & Rossion, 2016). This is done because the amplitude at any given frequency is considered as a combination of signal and noise (Heinrich et al., 2009). Finally, Z-scores were computed based on the grand-averaged amplitude spectrum for each condition, to assess the significance of the response at each stimulation frequency (base-10Hz, categorical change-2Hz) and harmonics (Liu-Shuang et al., 2014a; Lochy et al., 2015). Z-scores (*Z(x)= x-mean(noise)/SD (noise*) were considered significant if larger than 3.1 (p < 0.001, one-tailed, signal > noise)

In order to quantify the periodic response distributed across several harmonics, the baseline subtracted amplitudes of significant harmonics (excluding the base stimulation frequency) were summed for each participant, task, and condition (Retter & Rossion, 2016).

## Results

### Behavioural colour-change task

We first analysed accuracy and response times in the two tasks with paired t-tests. Due to recording failures, the data of 3 participants were lost. There was a non-significant trend for higher accuracy (hit rate) in the bars task (96.9%) than in the cross task (95.23%) [t(18)=1.999; p=.061]. However, false alarms were higher in the bars (5.04%) than in the cross task (3.02%) [t(18)=2.648; p=.016], which could be expected as only the bars task contained No-Go trials (e.g., colour-change on one side only). Finally, correct hits RT were longer for the bars (501.3ms) than the cross task (462.6ms) [t(18)=3.065; p=.007]. In the bars task, there was a non-significant trend for RT on false alarms (No-Go trials) to be shorter than RT on Go trials (457.1ms vs 501.3ms) [t(18)=1.532; p=.143]. In sum, these results indicate that both tasks were very well performed by the participants, with the bars task being slightly more demanding than the cross task.

### Categorical discrimination responses

Across tasks and conditions, averaged Z-scores were significant (z>3.1, p<.001) for discrimination responses from 2Hz up to 8Hz (i.e., 4 harmonics). Scalp topographies of the sum of harmonics show a larger word-selective response in the left hemisphere (Figure 2), as in previous studies (de Rosa et al., 2022; Lochy et al., 2015). Based on this, we selected *a priori* electrodes of interest in the LH (including 5 electrodes around P07) as well as their contralateral homologues to define LH and RH regions-of-interest (ROI) for further analysis.

**Figure 2.**
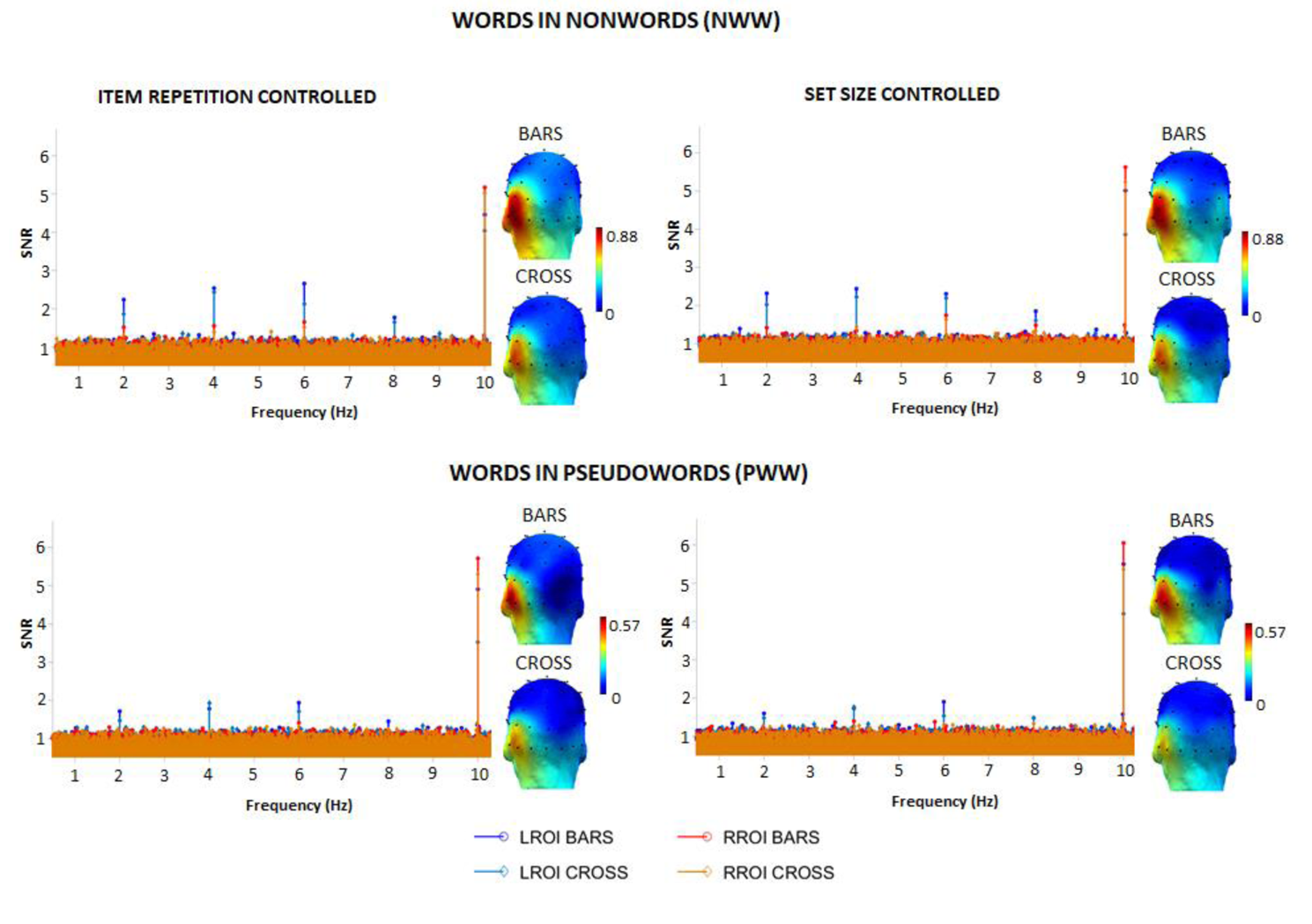
EEG spectra (SNR, baseline level = 1) and scalp topographies (scale in µV) of categorical discrimination responses for words (i.e., word-selective neural responses) among non-words (top row) or pseudowords (bottom row). The SNR is shown in left and right regions-of-interest (LROI/ RROI) and for each task (bars/cross). It reveals in each condition, task, and control type, clear discrimination responses for words at 2Hz and harmonics (significant up to 8Hz). The peak at 10Hz represents the base rate (general visual) response. While the type of control (set size or item repetition) did not influence response amplitudes, the orthogonal task induced stronger word discrimination responses when participants monitored two lateral vertical bars as a central cross.

The sum of baseline corrected amplitudes for the four significant harmonics were submitted to a 2 (*Tasks*: cross vs. bars) x 2 (*Discrimination level*: fine vs coarse-grained) x 2 (*Control Type*: set size vs. item repetition) x 2 (*Hemisphere*s: LH vs RH) repeated-measures ANOVA.

There was a main effect of *Hemisphere* [F(1,21)=24.219; p<.0001], with (about three times) larger responses in the LH (.473µV) than in the RH (.151µV) (Figure 2 and Figure 3). *Tasks* also significantly influenced response amplitudes [F(1,21)=5.206; p<.033), with larger EEG amplitudes in the bars (.342µV) than in the cross task (.282µV). There was a main effect of *Discrimination level* [F(1,21)=22.092; p<.0001), with stronger responses when words were inserted in NW (.411µV) than in PW (.213µV). In contrast, the factor *Control Type* was not significant [F(1,21)<1; p=.94; item repetition controlled: .311µV; set size controlled: .313µV).

**Figure 3.**
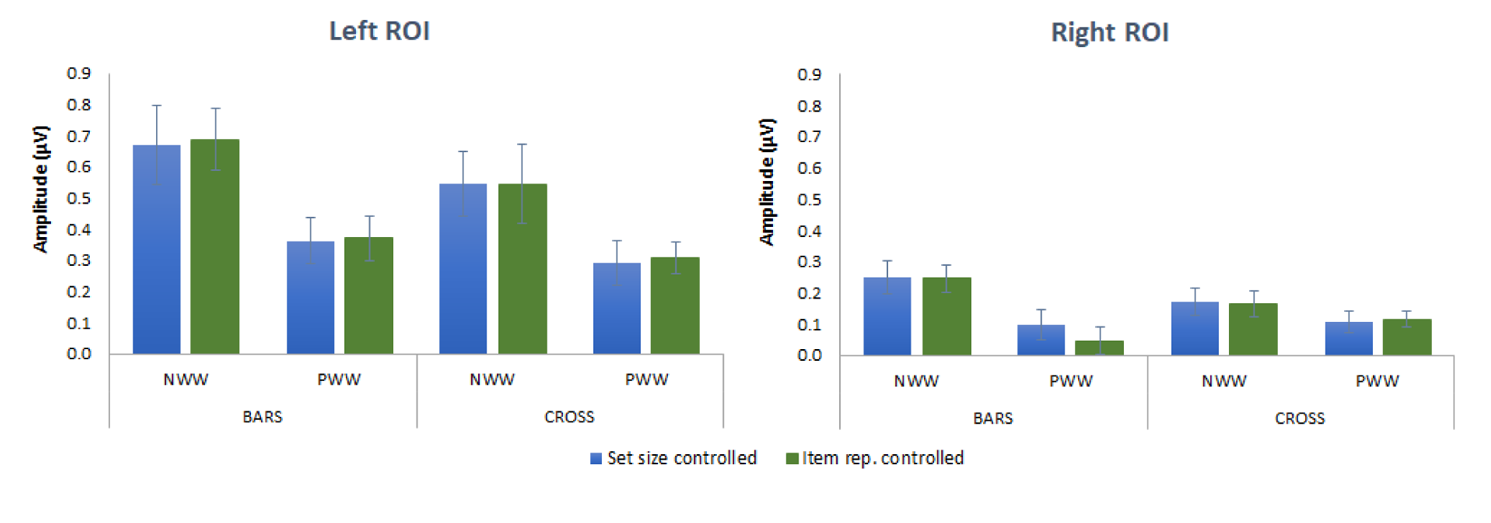
Word-selective EEG amplitudes (in µV, sum of harmonics), as a function of control type (blue: controlled set size; green: controlled item repetition), discrimination level (words among non-words: NWW; words among pseudowords: PWW) and task (bars, cross). The left panel represents the left ROI and the right panel the right ROI.

There were also several significant interactions modulating these effects. First, the interaction *Hemisphere* x *Task* [F(1,21)=7.266; p<.014) was due to an effect of *Task* in the LH [F(1,21)=7.687; p=.011] (cross: .42µV; bars: .52µV) but not in the RH (cross: .16; bars: .14µV). Second, there was an interaction between *Hemisphere* and *Discrimination level* [F(1,21)=6.456; p<.019]. Responses to words among NW (NWW) were higher than those among PW in each hemisphere (in the LH, t(21)=4.268; p<.0001; in the RH t(21)=3.203; p<.004), but the difference was stronger in the LH (.279µV) than in the RH (.116µV). The left dominance in response amplitude was present in each discrimination level as shown by paired t-tests contrasting hemispheres (for NWW, t(21)=4.455; p<.0001; and for PWW, t(21)=4.894; p<.0001). Finally, there was a trend for an interaction between *Task* and *Discrimination Level* [F(1,21)=4.196; p=.053), with the effect of task being marginally stronger for NWW sequences (bars: .465µV, cross: .357µV) than for PWW sequences (bars: .220µV, cross: .206µV). All other interactions were not significant (Fs<1).

### Base rate responses

Responses at the base rate (10Hz) were significant up to 4 harmonics (40Hz), when averaging across tasks and conditions, with the largest EEG amplitudes on posterior channels of the occipital medial region, as in previous studies (e.g., Lochy et al., 2015) (ranked as O2, Oz, Iz and O1 for the 4 first channels). We defined a medial-occipital region (MO-ROI) including Oz, Iz, O1 and O2 and we analysed the effect of *Tasks* (cross vs. bars) x *Discrimination level* (fine vs coarse-grained) x *Control Type* (item repetition vs set size) in a 2×2×2 repeated-measures ANOVA.

There were no significant main effects (all Fs <1) or interactions [*Task x Discrimination Level*, F(1,21)=2.472; p=.131; *Task x Item repetition* F(1,21)=1.329; p=.262; *Discrimination Level x Item repetition* F(1,21)=1.140; p=.298; *Task x Item repetition x Discrimination Level* F(1,21)=1.212; p=.283], suggesting that the overall general visual response did not differ across all these sequences showing strings of letters at 10Hz, whatever the task or nature of the contrast (Figure 4).

**Figure 4.**
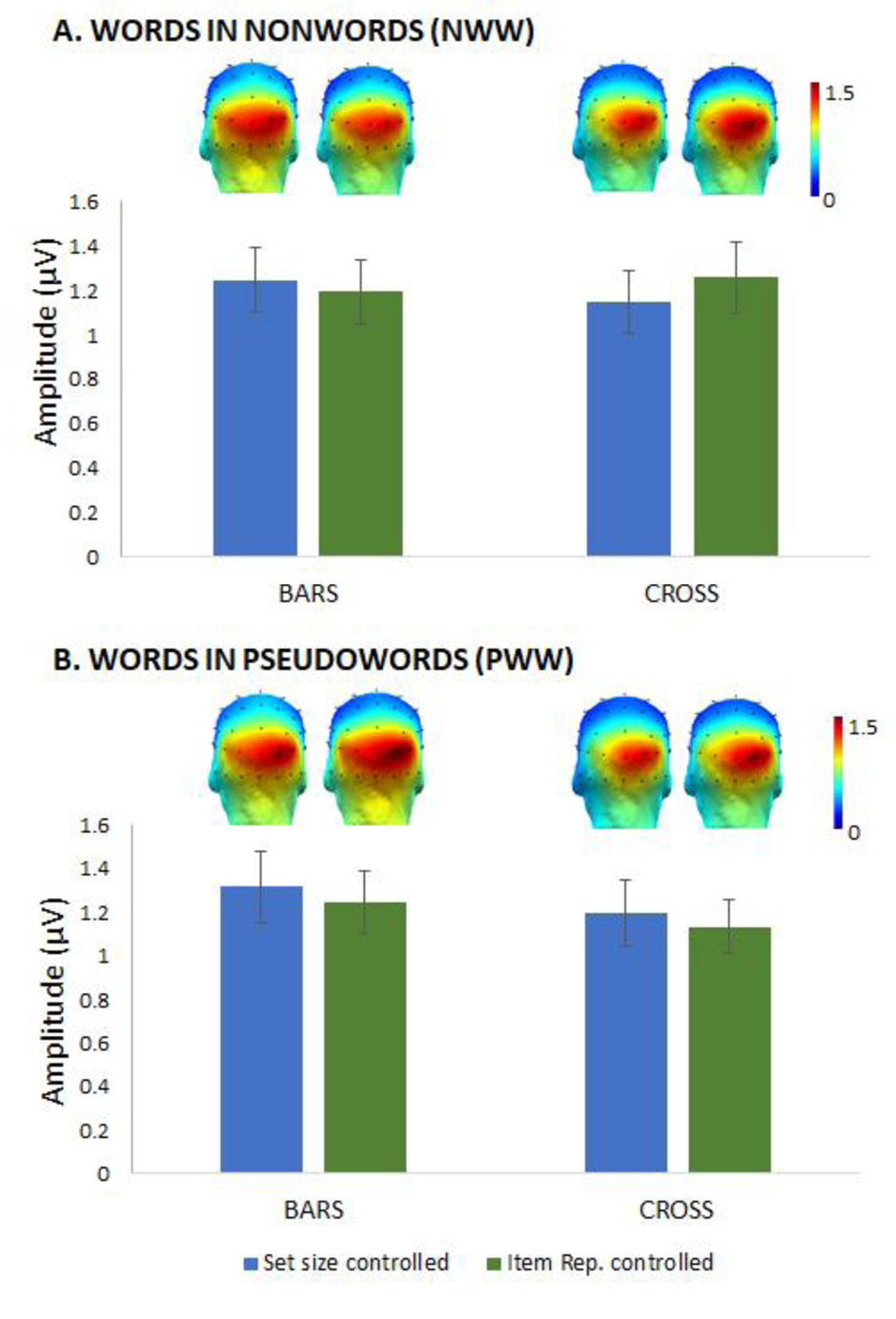
Base rate response amplitudes. (in µV) at a medial-occipital region of interest, as a function of *Discrimination Level* (**A.** words in nonwords, NWW; **B.** words in pseudowords, PWW), *Task* (cross vs. bars) and *Control Type* (item repetition vs. set size).

### Individual level analysis

Given the highly significant effects obtained when analysing data at the group-level, we also assessed whether the approach is powerful enough to evidence clear responses at the individual level (which would also be critical to know for potential clinical application with individual patients). For this, individual Z-scores were computed for each participant, discrimination level, control type, and task, in each ROI, on non-corrected summed amplitudes for significant harmonics as determined at the group level (see supplementary Figure S1 for display of individual Z-scores and Figure S2 with details in each cell of the design.).

Impressively, all of the 22 individual participants had a significant EEG response (z >1.64) for words among non-words in *at least* one ROI, task, and control type. 15 out of 22 (68%) had a significant response *in all conditions* (independently of ROI), and no individual showed no response at all. For the fine discrimination level, i.e., words among pseudowords, 20 out of 22 participants showed a significant response in *at least* one ROI, task, and control type. Among them, 10/22 (45%) showed a significant response *in all conditions*, and only two individuals did not show any significant response.

Considering the left ROI only, as most of the word-selective response concentrates in that region, and not taking into account the *Control Type* factor (as it does not influence responses at the group level), the sensitivity of the paradigm is extremely good as can be seen on Figure 5. For NWW, we were able to measure a significant response (set size or item repetition controlled) in 95% of participants, and for PWW, in 82% in the most sensitive bars task.

**Figure 5.**
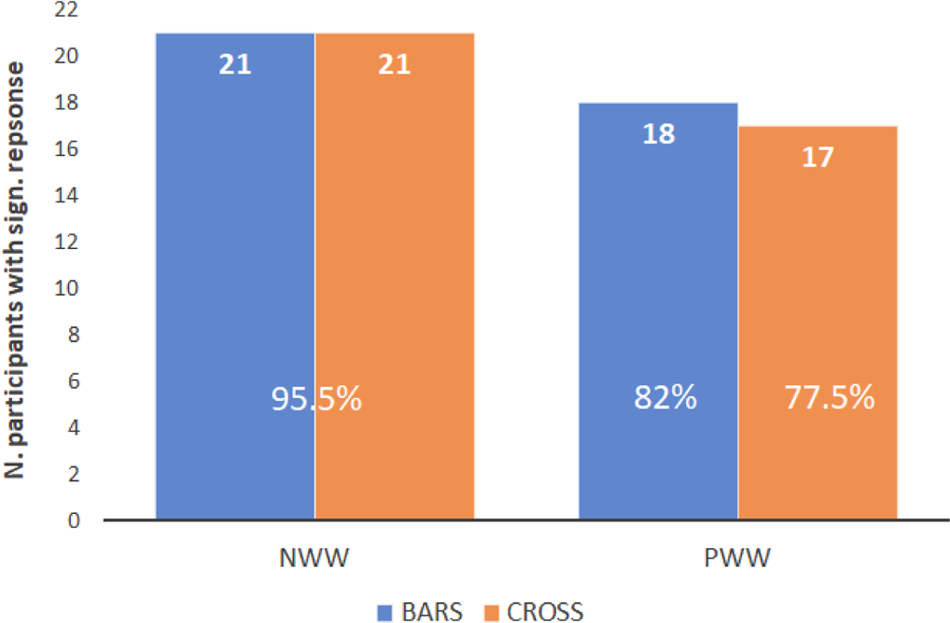
Number of participants (out of 22) showing a significant response in the left ROI per *discrimination coarseness condition* (NWW, PWW) and *task* (Bars, in blue and Cross in orange on the graph), in at least one of the *Control Types* conditions (item repetition or set size controlled).

We also computed the correlation across individuals between conditions (bars vs. cross, item number vs. repetition controlled; PW vs. NW) after calculating for each participant the average amplitude response in the LROI in each of the variables of interest.

Correlations were significant and high (Figure 6) between the two *Control Types* (item repetition vs set size: Spearman Rho=.810; p<.001); between cross and bars tasks (Rho=.816; p<.001); and slightly lower between the two discrimination levels (Rho=.740; p<.001).

**Figure 6.**
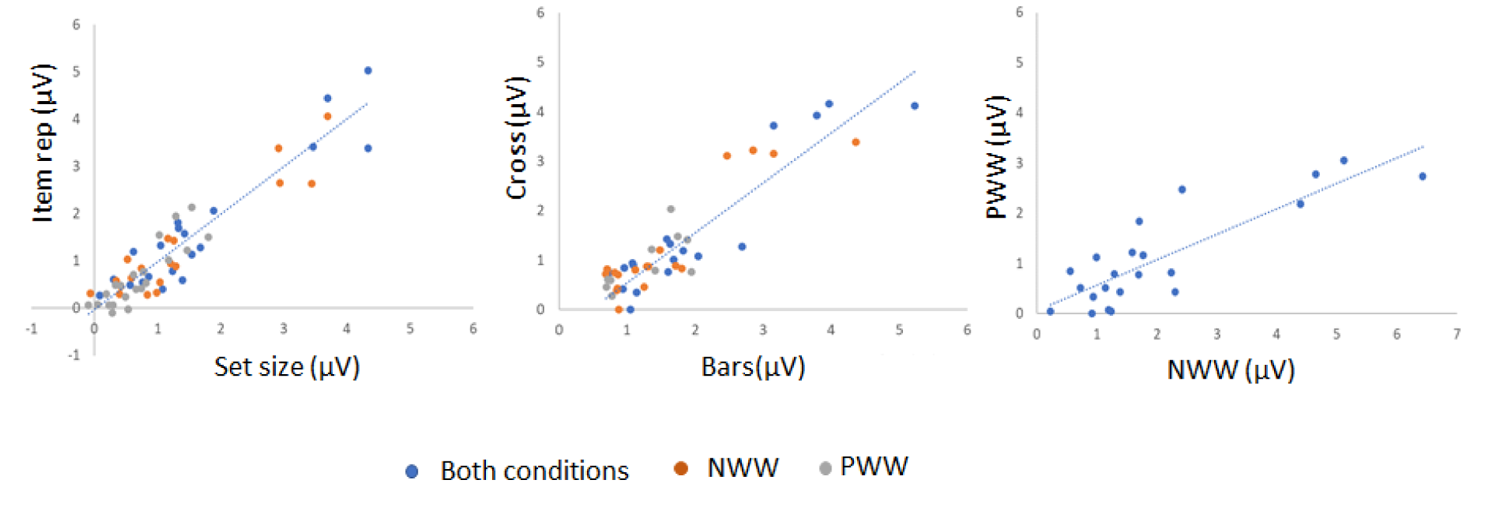
Scatter plots for individual word-selective responses illustrating correlations between: A. set size vs item repetition controlled; B. cross vs bars tasks; C. words in Pseudowords vs words in nonwords discrimination levels. In A & B, correlations were computed for both PWW and NWW conditions averaged, but individual data are also plotted for each condition separately (orange dots: NWW; grey dots: PWW).

Finally, given that the increase of discrimination responses with deployed attention could be due to a general arousal effect due to the bars task being slightly more demanding, we correlated the amplitude increase between cross and bars tasks (averaging Conditions and Control Types), to the RT increase between these two tasks. These variables did not correlate, considering only the left ROI (Spearman Rho=.091; p=.729), or both ROIs together (Spearman Rho=.23; p=.37).

## Discussion

Given the development and increasing use of an FPVS-oddball approach to study written letter and word-selective representations (since Barnes et al., 2021; de Rosa et al., 2022; Lochy et al., 2015, 2016, 2018; Lochy & Schiltz, 2019; van de Walle de Ghelcke et al., 2020, 2021; Wang et al., 2021), the present study explored in detail the nature of the word-selective response being measured. Specifically, we investigated both conceptual categorization of words (processed at linguistic levels), as well as more peripheral or design-related processes. For this, we used our standard FPVS-oddball-like paradigm in EEG and investigated the modulation of neural responses to words according to three key aspects of the paradigm: the discrimination coarseness (prelexical/ lexical), the relative item frequency (item repetition), and the use of a focused or deployed attentional task. Strengthening previous observations (Lochy et al., 2015), we found that the coarser discrimination contrast (prelexical) gives rise to the largest responses, but also that lexical responses are measurable at the individual participant level. Second, despite the high sensitivity of the paradigm, we found no evidence of an impact of item frequency on oddball responses. These two findings strongly support the interpretation of FPVS word-selective responses as stemming from psycholinguistic processing of words. Third, a deployed attentional task seems more optimal to use with the paradigm, as it positively influenced word discrimination in all conditions. Finally, we also demonstrate the sensitivity of the paradigm at the individual level. These findings are discussed below.

### Different levels of word-selectivity

With new stimuli and a larger sample of participants, we replicated a modulation of response amplitude as a function of discrimination coarseness, implemented by manipulating the wordlikeness of base stimuli (Lochy et al., 2015): word-selective responses were significantly larger when contrasted to non-words than pseudowords.

This finding is reminiscent of long-known list context effects on performance for visual word recognition at the behavioural level (Evans et al., 2012; Lupker & Pexman, 2010), accounted for within different theoretical frameworks (Grainger & Jacobs, 1996; Norris, 2006; Perry et al., 2007). In lexical decision for instance, the time to decide that an item is a word is shorter if inserted in non-wordlike as in wordlike lists. This can be explained at the cognitive level by the fact that the threshold for word recognition is modulated by the wordlikeness of nonwords (Lupker & Pexman, 2010). Each letter string activates lexical units containing shared letters/ spellings (i.e., neighbors) (Grainger & Jacobs, 1996). When presented with non-wordlike nonwords, which have none or very few orthographic neighbors, the overall level of activation in the lexicon is low and therefore the threshold of activation sufficient to recognize an item as a word can be modest (faster decision times). On the contrary, when the nonwords are very wordlike, then the overall level of activity in the lexicon is higher, and the threshold for word recognition needs to be higher (longer decision times). In our case, no explicit decision had to be made by participants, but we can still consider that in PWW sequences, upon each PW presentation (more wordlike), a set of neighbouring items is activated in the lexicon, while it is not the case in NWW sequences (less wordlike). In this context, when words are presented, they generate an overall lower difference of activation threshold in PWW by comparison to NWW sequences. However, it is important to emphasize that our task did not require any decision about lexicality, which would suggest that these effects stem from pre-decisional stages of processing. The simple viewing of words in different wordlikeness contexts was sufficient to trigger these effects, which thus supports the proposal that visual word recognition is unintentional and does not require explicit attention to the stimulus (Brown et al., 2002). However, one could also consider that a shallow processing of words, bypassing lexical processing, could be sufficient to trigger responses in NWW sequences. Indeed, statistical regularities in the combination of co-occurring letters are coded at the neural level (open-bigram model of word recognition, (Bouhali et al., 2014, 2019; Dehaene et al., 2005; Grainger & van Heuven, 2003; Mariol et al., 2008; Vinckier et al., 2007), and NW differed from words on crude prelexical/orthographic factors such as bigram frequency, or consonant-vowel structure (Table 1). As such, the paradigm does not allow us to ascertain if words were *also* processed at the lexical level when inserted in NW. This could be examined in future studies by contrasting responses to words vs. similarly valid orthographic forms like PW, both inserted in different NWW sequences. In this case, responses to PW can only reflect orthographic processing, while responses to words may be due to a combination of prelexical and lexical processes. In the PWW condition on the contrary, these pre-lexical factors were controlled, as the lists of W and PW did not differ in overall bigram frequency, orthographic neighbourhood, and consonant-vowel structure. We can thus safely interpret this word response as stemming from lexical levels (whole word-forms are recognized) or at semantic levels (only words have a meaning).

Regardless of the level of contrast, the left occipito-temporal scalp topography of word-selective activity is in agreement with the known brain regions implicated in reading and visual word recognition, such as the left VOTC (ventral occipito-temporal cortex) or VWFA (visual word form area) (Cattinelli et al., 2013; Dehaene et al., 2005; Schurz et al., 2014; Wandell & Le, 2017), and with word-selective intracerebral responses disclosed with this paradigm in the lateral portion left middle fusiform gyrus/occipito-temporal sulcus (Lochy et al., 2018). At the neural level, the index of differential processing measured by this paradigm is lower for PWW than NWW sequences. This presumably stems from the fact that populations of neurons responding to letter-strings are more similar, or partially overlapping, between PW and W, than between NW and W (Glezer et al., 2009). Indeed, an intracerebral FPVS study (Lochy et al., 2018) suggested that both fine-tuning to whole-words (e.g., lexical representations) and broad-tuning to plausible letter-strings (e.g., prelexical representations) are located in overlapping neural populations in the left fusiform gyrus (Glezer et al., 2009; Lochy et al., 2018). The sensitivity to wordlikeness of base stimuli suggests that both mechanisms were playing a role here. Broad-tuning to letter-strings could be sufficient to trigger strong word-selective response in the case of NW base stimuli, while fine-tuned neural mechanisms were necessary when words were presented in PWW base stimuli. This could be addressed in future studies using combined EEG-MEG and source estimation (e.g., Hauk et al., 2021).

### Statistical regularities do not contribute to word-selective responses

The FPVS-oddball paradigm is very sensitive for capturing the neural discrimination between a base and a deviant category of stimuli (for instance words as deviants among non-words or pseudowords as done here). It does, however, contain an inherent imbalance in exemplar repetition when, as often the case in such studies, set sizes of the two contrasted categories are equated. That is, given the importance of implicit statistical learning as a basic mechanism for extracting patterned regularities in the environment (Armstrong et al., 2017; Aslin, 2017; Christiansen, 2019; Frost et al., 2019), one could legitimately wonder if word-selective neural responses in previous studies attributed to their psycholinguistic characteristics only, mainly or partly reflected the detection of different frequency-of-occurrence of stimuli. Thus, one of the aims of our study was to evaluate if, and how much of, the discrimination response for words in the FPVS-oddball approach is at least partly due to detection of item repetition differences rather than detection of words as lexical items with specific properties by comparison to base stimuli.

If there were cumulative effects of statistical learning about regularities and orthographic/lexical processes, responses should have been larger when item repetition was imbalanced (set size controlled). However, contrary to this prediction, our results were crystal-clear: word-selective responses did not differ in amplitude or lateralization whether set size or item repetition were controlled. In addition, the correlation across individuals between these two conditions was very high (rho=.81), arguing for a qualitatively similar process underlying the word-selective response in both cases. Thus, there was no effect that could be attributed to statistical learning about exemplar repetition when using a deviant set of 30 items being presented less often than the base set of 30 items.

These observations are consistent with studies using this oddball-like paradigm in other domains. For instance, when contrasting (deviant) faces to (base) objects, the number of repetitions for exemplars in each category does not matter (Retter & Rossion, 2016). In this latter study, using a set of 46 faces and 248 objects images, the authors showed that the periodicity of deviant faces in 120s sequences presented at 12.5Hz, from 1 face every 5 objects to 1 every 11 objects (ratios face/objects: 1/5, 1/7, 1/9 and 1/11) had no influence whatsoever on the amplitude and spatial localization of the face-selective response. In another study, faces were presented at a fixed 1Hz rate in sequences varying in overall stimulation frequency, from 20Hz to 3Hz (Retter et al., 2020). In that case, the number of objects between face-deviants varied accordingly, i.e., faces were separated by 2 objects (3 Hz), or 5 (6 Hz), 11 (12 Hz) or even 19 (20 Hz). Again, the number of repetitions between deviants did not show any effect on amplitude of the face-selective response (Retter et al., 2020).

In contrast to these studies on faces and the present study on written words, we note that recent studies that found effects of statistical regularities, either by arbitrary grouping digits (Guillaume et al., 2020), or letters/characters-strings (de Rosa et al., 2022) used rather small sets of 4 or 8 items. Since statistical regularities must be easier to detect when the exact small set of stimuli is repeated, we think this factor could be key in statistical learning of novel groupings. If this is correct, future work should identify how many different items should be used in order to avoid these statistical learning effects. Since there are areas of research in which it is not possible to use many different items (e.g., in numerical cognition, when using Arabic digits as stimuli, there are maximum 10 elements (1 to 9, and 0)), future studies with FPVS-oddball like paradigms should be particularly careful, using changes in surface features of the stimuli to reduce/eliminate mere statistical regularity effects (see e.g., Marlair et al., 2022).

Interestingly, while we replicated the LH dominance in the scalp topography of word-selective responses, the study of de Rosa et al. (2022), identifying statistical learning effects with character strings found rather different scalp topographies: these responses were bilateral at posterior sites, without any effect of hemisphere. Furthermore, responses did not differ between conditions (words, pseudowords, non-words or artificial script), even though several more clusters were found for words. This posterior topography fits well with the visual networks that are activated by statistical learning tasks constituted of visual stimuli (Gavornik & Bear, 2014; Kok et al., 2012; Richter & de Lange, 2019). In contrast, our observation of a clear LH dominance is yet another argument in favour of interpreting the origin of the discrimination effect for words among NW or PW as being orthographic or lexical, rather than a mere detection of imbalance in item repetition.

### Deployed vs. focal attention

We found a relatively large effect of the attentional task, with larger word-selective responses in the deployed attention task than the focused attention task. The automatic word processing occurring in FPVS-EEG studies can therefore be modulated by an attentional manipulation, the same way Stroop effects are reduced by focusing spatial attention on a single letter of the word (e.g., Lachter et al., 2008; Stolz & Besner, 1999). The attention literature in general shows that too much focused attention may not be optimal, and that better performance in perceptual identification is obtained with an adequate allocation of attentional resources (Yeshurun & Carrasco, 1998). In written word recognition, a whole field of research investigated the effects of spatial cueing, aiming at assessing the consequences of processing words at an attended/unattended location, at understanding the spontaneous asymmetry of attention distribution to one or the other visual field, and at assessing whether word processing requires attention or is immune to attentional orientation (Franceschini et al., 2012; Ramamurthy et al., 2021; Stein, 2014; White et al., 2019). Also, studies have compared effects of spatial cueing on different types of written strings, reasoning that only the phonological route (hence, pseudoword identification) requires parsing into small units and therefore focused spatial attention (Ans et al., 1998; Perry et al., 2007); while recognizing whole words involves large grain-size processes, less affected by attentional manipulation. The results of these studies are inconsistent, some studies suggesting that the lexical status of the letter-strings (e.g., words or PW) does influence results (Auclair & Siéroff, 2002), while others report no difference in cueing effects as a function of the type of string (McCann et al., 1992), or even no cueing effect when stimuli are presented in central vision (Ducrot & Grainger, 2007). Most of these studies contrasted valid to invalid cueing (e.g., Posner’s paradigm) a situation which is far from what was done here. However, directly relevant for our findings, some studies also used a control or neutral cue that broadens the attention on both left and right spatial locations, similarly to what was done in the current study (Montani et al., 2014). In this study, contrasting PW, high-frequency words (HFW) and low-frequency words, the authors found a modulation of cueing effects by the lexical status and familiarity of the words. Recognition of HFWs was *facilitated* by the more global processing induced by the neutral cue, even more than the valid cue, presumably because this broad distribution of attention is the default mode for processing HFWs (Brand-D’abriesca & Lavie, 2007; Ghahghaei et al., 2013; Kliegl et al., 2006). This enhancement of word identification when attention is broadly distributed could explain our finding for greater discrimination responses when the task involved deploying visual attention over the two vertical bars to monitor.

Another non-negligeable factor lies in the visibility of all letters which is maintained with the deployed attention task, while the central cross could partly mask the central letter (depending on its identity and shape) when using stimuli of 5 letters as done here. This could reduce recognition processes for some of the words and trigger misperception of some of the PW (as words), therefore breaking the periodicity of the words perceived occurrence, and thereby, reducing the selective response at the deviant’s frequency.

### Individual sensitivity

Assessing the significance of responses at the individual level and the correlations between conditions gave rise to several interesting findings. First, the high correlation across individuals between responses to words when item repetition vs set size were controlled confirms that the lack of difference observed at the group level is not due to unstable/variable responses. Second, the high co-variation of amplitudes between the two attentional tasks suggests that the greater neural response with the bars task is due to a mere quantitative (deployment of spatial attention) and not a qualitative (e.g., increase in arousal) difference in word processing. This interpretation is reinforced by the lack of correlation between RT increase at the behavioural level between cross and bars tasks and the amplitude increase at the neural level between these two tasks. Finally, correlation was smaller but still significant between word-selective responses when inserted among NW vs PW, suggesting that these two processes of pre-lexical vs lexical access vary similarly across individuals.

Regarding the significance of responses at the individual level, in the coarser contrast (NWW), 100% of participants had at least one significant response for discriminating words, and more than 2/3^rd^ of them (68%) showed significant responses in all four conditions (2 Control Type x 2 Tasks). Results were more mitigated for lexical responses (PWW), where 2 participants never had any word-response (9%), while about half showed consistent responses in <u>all</u> conditions (45%). Nevertheless, when taking only the LROI (where the response concentrates) and the most sensitive attentional bars task, the rate of significant responses at the lexical level (words in PW) was 82% (18/22 participants), which is very high. It contrasts with a recent study using English words and pseudowords in a similar design (Barnes et al., 2021) where only 10% detection rate was found (testing 10 participants). We can only speculate on reasons for this discrepancy. First, we used higher variability than the latter study when generating pseudowords: we kept the syllabic and consonant-vowel structure matched item-wise, but we did not control for letter identity. In (Barnes et al., 2021), items were 4 letter-long, and there was only 1-letter difference between each of the four created PW and their matched words, and each time at the same first position (e.g., “BOOK”, wook, dook, gook, mook). Thus, each word and its 4 matched pseudowords activated the same set of orthographic neighbours in the lexicon. In the orthographic priming literature, unmasked pseudoword neighbour primes have been shown to play an inhibitory role in the word selection process during recognition (Burt, 2009; Carreiras et al., 1997; De Moor & Verguts, 2006; Grainger & Segui, 1990; Versace, 1998). Thus, there might be an impact of repeatedly activating the same orthographic neighbours, acting as competitor items in the lexicon on further recognition processes of target words. Second, language was different. While we tested French-speaking participants, (Barnes et al., 2021) studied English-speaking participants. These two languages vary in several orthographic properties, such as transparency, or the number of orthographic neighbours. English words, for instance, have a greater orthographic neighbourhood size than French words overall, but the difference is even more striking when considering the stimuli used in the two respective experiments: the mean number of orthographic neighbours *N* is 19 for 4-letter English words, while it is 6 only for 5-letters French words (Marian et al., 2012). Neighbourhood size has been shown inconsistently to play a facilitatory (Grainger, 1992; Perea, 2015) or inhibitory (Perea et al., 2008; Pollatsek et al., 1999; Van Heuven et al., 1998; Whitney & Lavidor, 2005) role on word recognition, although the picture is further complicated by both frequency effects and task demands (Grainger & Segui, 1990). In any case, the questions raised by result incongruencies in FPVS studies using words show that it is crucial to design new experiments to understand if the lack of effects observed in (Barnes et al., 2021), in contrast to significant findings here, are due to methodological choices or if language properties as evoked above may play an intrinsic role.

### Summary and conclusions

In conclusion, the current study brings several important new findings that help strengthening and understanding the nature of word-selective neural responses obtained in FPVS-EEG studies. First, at least when using a relatively large stimulus set, the FPVS-oddball paradigm does measure word-selective responses because of their orthographic or lexical properties and not because of statistical (learning of) (ir)regularities during the stimulation sequences. This is evidenced both by the lack of influence of controlling item repetition, by the LH topography of the response, and by the modulation of the response by the discrimination level (wordlikeness of base stimuli). Second, the orthogonal task performed by our participants has an important impact, with greater discrimination responses when attention is deployed on both sides of the letters-strings rather than being focused in the centre. Third, word-selective neural responses are more clearly identified when the discrimination is coarser (words among nonwords) presumably reflecting a mixture of pre-lexical and lexical processes. In the former case, individual responses in the LROI are significant in 95% of the cases, while they are significant in 82% of the cases when they are discriminated from PW, reinforcing previous findings in French-speaking participants (Lochy et al., 2015).

## Acknowledgements

The project was partly funded by a *Lorraine Université d’Excellence* (LUE) grant to foster international collaborations between Université de Lorraine and University of Luxembourg (UL_IRP 2022). The first author is supported by the Fonds National de la Recherche du Luxembourg (FNR-CORE C21/SC/16241557/READINGBRAIN).

## References

Ans, B., Carbonnel, S., & Valdois, S. (1998). A connectionist multiple-trace memory model for polysyllabic word reading. Psychological Review, 105(4), 678.

Arciuli, J., & Simpson, I. C. (2012). Statistical Learning Is Related to Reading Ability in Children and Adults. Cognitive Science, 36(2), 286–304. https://doi.org/10.1111/j.1551-6709.2011.01200.x

Aristei, S., Lochy, A., Rossion, B., & Schiltz, C. (2017). A task independent brain signature of multilingual early visual word recognition. Conference of the European Society for Cognitive Psychology (ESCoP).

Armstrong, B. C., Frost, R., & Christiansen, M. H. (2017). The long road of statistical learning research: past, present and future. Philosophical Transactions of the Royal Society of London. Series B, Biological Sciences, 372(1711), 1–4. https://doi.org/10.1098/rstb.2016.0047

Aslin, R. N. (2017). Statistical learning: A powerful mechanism that operates by mere exposure. Wiley Interdisciplinary Reviews: Cognitive Science, 8(1–2), e1373.

Auclair, L., & Siéroff, E. (2002). Attentional cueing effect in the identification of words and pseudowords of different length. Quarterly Journal of Experimental Psychology Section A: Human Experimental Psychology, 55(2), 445–463. https://doi.org/10.1080/02724980143000415

Barnes, L., Petit, S., Badcock, N. A., Whyte, C. J., & Woolgar, A. (2021). Word Detection in Individual Subjects Is Difficult to Probe With Fast Periodic Visual Stimulation. Frontiers in Neuroscience, 15(March). https://doi.org/10.3389/fnins.2021.602798

Bouhali, F., Bézagu, Z., Dehaene, S., & Cohen, L. (2019). A mesial-to-lateral dissociation for orthographic processing in the visual cortex. Proceedings of the National Academy of Sciences of the United States of America, 116(43), 21936–21946. https://doi.org/10.1073/pnas.1904184116

Bouhali, F., Thiebaut de Schotten, M., Pinel, P., Poupon, C., Mangin, J.-F., Dehaene, S., & Cohen, L. (2014). Anatomical connections of the visual word form area. The Journal of Neuroscience: The Official Journal of the Society for Neuroscience, 34(46), 15402–15414. https://doi.org/10.1523/JNEUROSCI.4918-13.2014

Brand-D’abriesca, M., & Lavie, N. (2007). Distractor effects during processing of words under load. Psychonomic Bulletin & Review, 14(6), 1153–1157. https://doi.org/10.3758/BF03193105

Brown, T. L., Gore, C. L., & Carr, T. H. (2002). Visual attention and word recognition in Stroop color naming: Is word recognition “automatic?” Journal of Experimental Psychology: General, 131(2). https://doi.org/10.1037//0096-3445.131.2.220

Burt, J. S. (2009). Identifiable orthographically similar word primes interfere in visual word identification. Journal of Memory and Language, 61(3), 259–284. https://doi.org/10.1016/j.jml.2009.06.003

Carreiras, M., Perea, M., & Grainger, J. (1997). Effects of orthographic neighborhood in visual word recognition: Cross-task comparisons. Journal of Experimental Psychology: Learning Memory and Cognition, 23(4), 857–871. https://doi.org/10.1037/0278-7393.23.4.857

Cattinelli, I., Borghese, N. A., Gallucci, M., & Paulesu, E. (2013). Reading the reading brain: A new meta-analysis of functional imaging data on reading. Journal of Neurolinguistics, 26(1), 214– 238. https://doi.org/10.1016/j.jneuroling.2012.08.001

Christiansen, M. H. (2019). Implicit Statistical Learning: A Tale of Two Literatures. Topics in Cognitive Science, 11(3), 468–481. https://doi.org/10.1111/tops.12332

de Heering, A., & Rossion, B. (2015). Rapid categorization of natural face images in the infant right hemisphere. ELife, 4, 1–14. https://doi.org/10.7554/eLife.06564

De Moor, W., & Verguts, T. (2006). When are neighbours hostile? Inhibitory neighbour effects in visual word recognition. Psychological Research, 70(5), 359–366. https://doi.org/10.1007/S00426-005-0225-X

de Rosa, M., Ktori, M., Vidal, Y., Bottini, R., & Crepaldi, D. (2022). Frequency-based neural discrimination in fast periodic visual stimulation. Cortex, 148(January), 193–203. https://doi.org/10.1016/j.cortex.2022.01.005

Dehaene, S., Cohen, L., Sigman, M., & Vinckier, F. (2005). The neural code for written words: a proposal. Trends in Cognitive Sciences, 9(7), 335–341. https://doi.org/10.1016/j.tics.2005.05.004

Dienes, Z., Broadbent, D., & Berry, D. (1991). Implicit and Explicit Knowledge Bases in Artificial Grammar Learning. *Journal of Experimental Psychology: Learning*, Memory, and Cognition, 17(5), 875–887. https://doi.org/10.1037/0278-7393.17.5.875

Ducrot, S., & Grainger, J. (2007). Deployment of spatial attention to words in central and peripheral vision. Perception and Psychophysics, 69(4), 578–590. https://doi.org/10.3758/BF03193915

Duyck, W., Desmet, T., Verbeke, L. P. C., & Brysbaert, M. (2004). WordGen: A tool for word selection and nonword generation in Dutch, English, German, and French. Behavior Research Methods, Instruments, & Computers, 36(3), 488–499. https://doi.org/10.3758/BF03195595

Eriksen, C. W., & St James, J. D. (1986). Visual attention within and around the field of focal attention: A zoom lens model. In Perception & Psychophysics (Vol. 40, Issue 4).

Evans, G. a L., Lambon Ralph, M. a, & Woollams, A. M. (2012). What’s in a word? A parametric study of semantic influences on visual word recognition. Psychonomic Bulletin & Review, 19(2), 325– 331. https://doi.org/10.3758/s13423-011-0213-7

Franceschini, S., Gori, S., Ruffino, M., Pedrolli, K., & Facoetti, A. (2012). A causal link between visual spatial attention and reading acquisition. Current Biology, 22(9), 814–819. https://doi.org/10.1016/j.cub.2012.03.013

Frost, R., Armstrong, B. C., & Christiansen, M. H. (2019). Statistical learning research: A critical review and possible new directions. Psychological Bulletin, 145(12), 1128–1153. https://doi.org/10.1037/bul0000210

Gavornik, J. P., & Bear, M. F. (2014). Learned spatiotemporal sequence recognition and prediction in primary visual cortex. Nature Publishing Group, 17(5). https://doi.org/10.1038/nn.3683

Ghahghaei, S., Linnell, K. J., Fischer, M. H., Dubey, A., & Davis, R. (2013). Effects of load on the time course of attentional engagement, disengagement, and orienting in reading. Quarterly Journal of Experimental Psychology, 66(3), 453–470. https://doi.org/10.1080/17470218.2011.635795

Glezer, L. S., Jiang, X., & Riesenhuber, M. (2009). Evidence for highly selective neuronal tuning to whole words in the “visual word form area”. Neuron, 62(2), 199–204. https://doi.org/10.1016/j.neuron.2009.03.017

Grainger, J. (1992). Chapter 7 Orthographic Neighborhoods and Visual Word Recognition. In R. Frost & L. B. T.-A. in P. Katz (Eds.), Orthography, Phonology, Morphology, and Meaning (Vol. 94, pp. 131–146). North-Holland. https://doi.org/10.1016/S0166-4115(08)62792-2

Grainger, J., & Jacobs, A. M. (1996). Orthographic Processing in Visual Word Recognition: A Multiple Read-Out Model. Psychological Review, 103(3), 518–565. http://www.scopus.com/inward/record.url?eid=2-s2.0-0030195303&partnerID=tZOtx3y1

Grainger, J., & Segui, J. (1990). Neighborhood frequency effects in visual word recognition: A comparison of lexical decision and masked identification latencies. Perception & Psychophysics, 47(2), 191–198. https://doi.org/10.3758/BF03205983

Grainger, J., & van Heuven, W. J. B. (2003). Modeling letter position coding in printed word perception. In The mental lexicon.

Guillaume, M., Mejias, S., Rossion, B., Dzhelyova, M., & Schiltz, C. (2018). A rapid, objective and implicit measure of visual quantity discrimination. Neuropsychologia, 111(July 2017), 180–189. https://doi.org/10.1016/j.neuropsychologia.2018.01.044

Guillaume, M., Poncin, A., Schiltz, C., & van Rinsveld, A. (2020). Measuring spontaneous and automatic processing of magnitude and parity information of Arabic digits by frequency-tagging EEG. Scientific Reports, 10(1), 1–11. https://doi.org/10.1038/s41598-020-79404-w

Hauk, O., Rice, G. E., Volfart, A., Magnabosco, F., Ralph, M. A. L., & Rossion, B. (2021). Face-selective responses in combined EEG/MEG recordings with fast periodic visual stimulation (FPVS). NeuroImage, 242. https://doi.org/10.1016/j.neuroimage.2021.118460

Heinrich, S. P., Mell, D., & Bach, M. (2009). Frequency-domain analysis of fast oddball responses to visual stimuli: a feasibility study. International Journal of Psychophysiology: Official Journal of the International Organization of Psychophysiology, 73(3), 287–293. https://doi.org/10.1016/j.ijpsycho.2009.04.011

Kidd, E., & Arciuli, J. (2016). Individual Differences in Statistical Learning Predict Children’s Comprehension of Syntax. 87(1), 184–193. https://doi.org/10.1111/cdev.12461

Kliegl, R., Nuthmann, A., & Engbert, R. (2006). Tracking the mind during reading: The influence of past, present, and future words on fixation durations. Journal of Experimental Psychology: General, 135(1), 12–35. https://doi.org/10.1037/0096-3445.135.1.12

Kok, P., Jehee, J. F. M., & de Lange, F. P. (2012). Less Is More: Expectation Sharpens Representations in the Primary Visual Cortex. Neuron, 75(2), 265–270. https://doi.org/10.1016/j.neuron.2012.04.034

Lachter, J., Ruthruff, E., Lien, M. C., & McCann, R. S. (2008). Is attention needed for word identification? Evidence from the Stroop paradigm. Psychonomic Bulletin and Review, 15(5), 950–955. https://doi.org/10.3758/PBR.15.5.950

Liu-Shuang, J., Norcia, A. M., & Rossion, B. (2014a). An objective index of individual face discrimination in the right occipito-temporal cortex by means of fast periodic oddball stimulation. Neuropsychologia, 52, 57–72. https://doi.org/10.1016/j.neuropsychologia.2013.10.022

Liu-Shuang, J., Norcia, A. M., & Rossion, B. (2014b). An objective index of individual face discrimination in the right occipito-temporal cortex by means of fast periodic oddball stimulation. Neuropsychologia, 52, 57–72. https://doi.org/10.1016/j.neuropsychologia.2013.10.022

Lochy, A., de Heering, A., & Rossion, B. (2019). The non-linear development of the right hemispheric specialization for human face perception. Neuropsychologia, 126, 10–19. https://doi.org/10.1016/j.neuropsychologia.2017.06.029

Lochy, A., Jacques, C., Maillard, L., Colnat-Coulbois, S., Rossion, B., & Jonas, J. (2018). Selective visual representation of letters and words in the left ventral occipito-temporal cortex with intracerebral recordings. Proceedings of the National Academy of Sciences of the United States of America, 115(32), E7595–E7604. https://doi.org/10.1073/pnas.1718987115

Lochy, A., & Schiltz, C. (2019). Lateralized Neural Responses to Letters and Digits in First Graders. Child Development, 90(6), 1866–1874. https://doi.org/10.1111/cdev.13337

Lochy, A., van Belle, G., & Rossion, B. (2015). A robust index of lexical representation in the left occipito-temporal cortex as evidenced by EEG responses to fast periodic visual stimulation. Neuropsychologia, 66, 18–31. https://doi.org/10.1016/j.neuropsychologia.2014.11.007

Lochy, A., van Reybroeck, M., & Rossion, B. (2016). Left cortical specialization for visual letter strings predicts rudimentary knowledge of letter-sound association in preschoolers. Proceedings of the National Academy of Sciences of the United States of America, 113(30), 8544–8549. https://doi.org/10.1073/pnas.1520366113

Lupker, S. J., & Pexman, P. M. (2010). Making things difficult in lexical decision: the impact of pseudohomophones and transposed-letter nonwords on frequency and semantic priming effects. Journal of Experimental Psychology. Learning, Memory, and Cognition, 36(5), 1267– 1289. https://doi.org/10.1037/a0020125

Marian, V., Bartolotti, J., Chabal, S., & Shook, A. (2012). Clearpond: Cross-linguistic easy-access resource for phonological and orthographic neighborhood densities. PLoS ONE, 7(8). https://doi.org/10.1371/journal.pone.0043230

Marinova, M., Georges, C., Guillaume, M., Reynvoet, B., Schiltz, C., & van Rinsveld, A. (2021). Automatic integration of numerical formats examined with frequency-tagged EEG. Scientific Reports, 11(1), 1–12. https://doi.org/10.1038/s41598-021-00738-0

Mariol, M., Jacques, C., Schelstraete, M. A., & Rossion, B. (2008). The speed of orthographic processing during lexical decision: Electrophysiological evidence for independent coding of letter identity and letter position in visual word recognition. Journal of Cognitive Neuroscience, 20(7). https://doi.org/10.1162/jocn.2008.20088

Marlair, C., Crollen, V., & Lochy, A. (2022). A shared numerical magnitude representation evidenced by the distance effect in frequency-tagging EEG. Scientific Reports, 12(1). https://doi.org/10.1038/s41598-022-18811-7

Marlair, C., Lochy, A., Buyle, M., Schiltz, C., & Crollen, V. (2021). Canonical representations of fingers and dots trigger an automatic activation of number semantics: an EEG study on 10-year-old children. Neuropsychologia, 157(August 2020). https://doi.org/10.1016/j.neuropsychologia.2021.107874

McCann, R. S., Folk, C. L., & Johnston, J. C. (1992). The Role of Spatial Attention in Visual Word Processing. Journal of Experimental Psychology: Human Perception and Performance, 18(4), 1015–1029. https://doi.org/10.1037/0096-1523.18.4.1015

Misyak, J. B., & Christiansen, M. H. (2012). Statistical Learning and Language: An Individual Differences Study. Language Learning, 62(1), 302–331. https://doi.org/10.1111/j.1467-9922.2010.00626.x

Misyak, J. B., Christiansen, M. H., & Tomblin, J. B. (2010). On-line individual differences in statistical learning predict language processing. Frontiers in Psychology, 1(SEP), 1–9. https://doi.org/10.3389/fpsyg.2010.00031

Montani, V., Facoetti, A., & Zorzi, M. (2014). Spatial attention in written word perception. Frontiers in Human Neuroscience, 8(February), 42. https://doi.org/10.3389/fnhum.2014.00042

Nazir, T. A., Ben-Boutayab, N., Decoppet, N., Deutsch, A., & Frost, R. (2004). Reading habits, perceptual learning, and recognition of printed words. Brain and Language, 88(3), 294–311. https://doi.org/10.1016/S0093-934X(03)00168-8

Norris, D. (2006). The Bayesian reader: explaining word recognition as an optimal Bayesian decision process. Psychological Review, 113(2), 327–357. https://doi.org/10.1037/0033-295X.113.2.327

Nurdal, V., Fairchild, G., & Stothart, G. (2021). The effect of repetition priming on implicit recognition memory as measured by Fast Periodic Visual Stimulation and EEG. International Journal of Psychophysiology, 161(December 2020), 44–52. https://doi.org/10.1016/j.ijpsycho.2021.01.009

Perea, M. (2015). Neighborhood effects in visual word recognition and reading. In The Oxford handbook of reading. (pp. 76–87). Oxford University Press.

Perea, M., Acha, J., & Fraga, I. (2008). Lexical competition is enhanced in the left hemisphere: Evidence from different types of orthographic neighbors. Brain and Language, 105(3), 199– 210. https://doi.org/10.1016/j.bandl.2007.08.005

Perry, C., Ziegler, J. C., & Zorzi, M. (2007). Nested incremental modeling in the development of computational theories: The CDP+ model of reading aloud. Psychological Review, 114(2), 273– 315. https://doi.org/10.1037/0033-295X.114.2.273

Peykarjou, S., Hoehl, S., Pauen, S., & Rossion, B. (2017). Rapid Categorization of Human and Ape Faces in 9-Month-Old Infants Revealed by Fast Periodic Visual Stimulation. Scientific Reports, 7(1), 1–12. https://doi.org/10.1038/s41598-017-12760-2

Pollatsek, A., Perea, M., & Binder, K. S. (1999). The effects of “neighborhood size” in reading and lexical decision. Journal of Experimental Psychology: Human Perception and Performance, 25(4), 1142–1158. https://doi.org/10.1037/0096-1523.25.4.1142

Posner, M. I. (1988). Structures and function of selective attention. In Clinical neuropsychology and brain function: Research, measurement, and practice. (pp. 173–202). American Psychological Association. https://doi.org/10.1037/10063-005

Ramamurthy, M., White, A. L., Chou, C., & Yeatman, J. D. (2021). Spatial attention in encoding letter combinations. Scientific Reports, 11(1), 1–12. https://doi.org/10.1038/s41598-021-03558-4

Rayner, K. (1998). Eye movements in reading and information processing: 20 years of research. Psychological Bulletin, 124(3), 372.

Retter, T. L., Jiang, F., Webster, M. A., & Rossion, B. (2020). All-or-none face categorization in the human brain. NeuroImage, 213(February), 116685. https://doi.org/10.1016/j.neuroimage.2020.116685

Retter, T. L., & Rossion, B. (2016). Uncovering the neural magnitude and spatio-temporal dynamics of natural image categorization in a fast visual stream. Neuropsychologia, 91, 9–28. https://doi.org/10.1016/j.neuropsychologia.2016.07.028

Richter, D., & de Lange, F. P. (2019). Statistical learning attenuates visual activity only for attended stimuli. ELife, 8, 1–27. https://doi.org/10.7554/eLife.47869

Rossion, B., & Boremanse, A. (2011). Robust sensitivity to facial identity in the right human occipito-temporal cortex as revealed by steady-state visual-evoked potentials. Journal of Vision, 11(2), 1–21. https://doi.org/10.1167/11.2.16

Rossion, B., Jacques, C., & Jonas, J. (2018). Mapping face categorization in the human ventral occipitotemporal cortex with direct neural intracranial recordings. Annals of the New York Academy of Sciences. https://doi.org/10.1111/nyas.13596

Rossion, B., Prieto, E. A., Boremanse, A., Kuefner, D., & Van Belle, G. (2012). A steady-state visual evoked potential approach to individual face perception: effect of inversion, contrast-reversal and temporal dynamics. NeuroImage, 63(3), 1585–1600. https://doi.org/10.1016/j.neuroimage.2012.08.033

Rossion, B., Retter, T. L., & Liu-Shuang, J. (2020). Understanding human individuation of unfamiliar faces with oddball fast periodic visual stimulation and electroencephalography. European Journal of Neuroscience, 52(10), 4283–4344. https://doi.org/10.1111/ejn.14865

Saffran, J. R., Aslin, R. N., & Newport, E. L. (1996). Statistical Learning by 8-Month-Old Infants. Science, 274(5294), 1926–1928. https://doi.org/10.1126/SCIENCE.274.5294.1926

Saffran, J. R., Newport, E. L., & Aslin, R. N. (1996). Word segmentation: The role of distributional cues. Journal of Memory and Language, 35(4), 606–621. https://doi.org/10.1006/JMLA.1996.0032

Schurz, M., Wimmer, H., Richlan, F., Ludersdorfer, P., Klackl, J., & Kronbichler, M. (2014). Resting-State and Task-Based Functional Brain Connectivity in Developmental Dyslexia. *Cerebral Cortex*, June 2015. https://doi.org/10.1093/cercor/bhu184

Shafto, C. L., Conway, C. M., Field, S. L., & Houston, D. M. (2012). Visual Sequence Learning in Infancy: Domain-General and Domain-Specific Associations With Language. Infancy, 17(3), 247– 271. https://doi.org/10.1111/j.1532-7078.2011.00085.x

Srinivasan, R., Russell, D. P., Edelman, G. M., & Tononi, G. (1999). Increased synchronization of neuromagnetic responses during conscious perception. Journal of Neuroscience, 19(13), 5435– 5448.

Stein, J. (2014). Dyslexia: the Role of Vision and Visual Attention. Current Developmental Disorders Reports, 1(4), 267–280. https://doi.org/10.1007/s40474-014-0030-6

Stolz, J. A., & Besner, D. (1999). On the myth of automatic semantic activation in reading. Current Directions in Psychological Science, 8(2), 61–65. https://doi.org/10.1111/1467-8721.00015

Stothart, G., Quadflieg, S., & Milton, A. (2017). A fast and implicit measure of semantic categorisation using steady state visual evoked potentials. Neuropsychologia, 102(December 2016), 11–18. https://doi.org/10.1016/j.neuropsychologia.2017.05.025

Turatto, M., Benso, F., Facoetti, A., Galfano, G., Mascetti, G., & Umilta, C. (2000). Automatic and voluntary focusing of attention. Perception & Psychophysics, 62, 935–952.

van de Walle de Ghelcke, A., Rossion, B., Schiltz, C., & Lochy, A. (2020). Impact of Learning to Read in a Mixed Approach on Neural Tuning to Words in Beginning Readers. Frontiers in Psychology, 10(January), 1–15. https://doi.org/10.3389/fpsyg.2019.03043

van de Walle de Ghelcke, A., Rossion, B., Schiltz, C., & Lochy, A. (2021). Developmental changes in neural letter-selectivity: A 1-year follow-up of beginning readers. Developmental Science, 24(1), 1–17. https://doi.org/10.1111/desc.12999

Van Heuven, W. J. B., Dijkstra, T., & Grainger, J. (1998). Orthographic Neighborhood Effects in Bilingual Word Recognition. Journal of Memory and Language, 39(3), 458–483. https://doi.org/10.1006/jmla.1998.2584

Versace, R. (1998). Frequency and prime duration effects on repetition priming and orthographic priming with words and pseudowords. Cahiers de Psychologie Cognitive/Current Psychology of Cognition, 17(3), 533–554.

Vettori, S., Dzhelyova, M., van der Donck, S., Jacques, C., Steyaert, J., Rossion, B., & Boets, B. (2019). Reduced neural sensitivity to rapid individual face discrimination in autism spectrum disorder. NeuroImage: Clinical, 21(November), 101613. https://doi.org/10.1016/j.nicl.2018.101613

Vinckier, F., Dehaene, S., Jobert, A., Dubus, J. P., Sigman, M., & Cohen, L. (2007). Hierarchical Coding of Letter Strings in the Ventral Stream: Dissecting the Inner Organization of the Visual Word-Form System. Neuron, 55(1), 143–156. https://doi.org/10.1016/j.neuron.2007.05.031

Volfart, A., Rice, G. E., Lambon, M. A., & Rossion, B. (2021). Implicit, automatic semantic word categorisation in the left occipito-temporal cortex as revealed by fast periodic visual stimulation. NeuroImage, 238(February), 118228. https://doi.org/10.1016/j.neuroimage.2021.118228

Wandell, B. A., & Le, R. K. (2017). Diagnosing the Neural Circuitry of Reading. Neuron, 96(2), 298– 311. https://doi.org/10.1016/j.neuron.2017.08.007

Wang, F., Kaneshiro, B., Strauber, C. B., Hasak, L., Nguyen, Q. T. H., Yakovleva, A., Vildavski, V. Y., Norcia, A. M., & McCandliss, B. D. (2021). Distinct neural sources underlying visual word form processing as revealed by steady state visual evoked potentials (SSVEP). Scientific Reports, 11(1), 1–15. https://doi.org/10.1038/s41598-021-95627-x

White, A. L., Boynton, G. M., & Yeatman, J. D. (2019). The link between reading ability and visual spatial attention across development. Cortex, 121, 44–59. https://doi.org/10.1016/j.cortex.2019.08.011

Whitney, C., & Lavidor, M. (2005). Facilitative orthographic neighborhood effects: The SERIOL model account. Cognitive Psychology, 51(3), 179–213. https://doi.org/10.1016/j.cogpsych.2005.07.001

Yan, X., Goffaux, V., & Rossion, B. (2022). Coarse-to-Fine(r) Automatic Familiar Face Recognition in the Human Brain. Cerebral Cortex, 32(8), 1560–1573. https://doi.org/10.1093/cercor/bhab238

Yeshurun, Y., & Carrasco, M. (1998). Attention improves or impairs visual performance by enhancing spatial resolution. Nature, 396(6706), 72–75. https://doi.org/10.1038/23936

