## supplementary material for "Linguistic and attentional factors – not statistical regularities – contribute to word-selective neural responses with FPVS-oddball paradigms"

**Table S1: stimuli**. List of 30 experimental words and their consonant-vowel structure, 4 lists of pseudowords (n=120) matched item-wise on CV structure, 4 lists of nonwords (n=120) matched at the letter-level to pseudowords. For mean characteristics at lexical and orthographic level, see text (Table 1).

| Words | CV structure | PW list1 | PW list2 | PW list3 | PW list4 | NW list1 | NW list2 | NW list3 | NW list4 |
|---|---|---|---|---|---|---|---|---|---|
| chose | CCVCV | crase | chire | brige | glite | rcesa | rcohi | rgebi | tgeli |
| chute | CCVCV | vrile | trute | flabe | glape | rvile | tteru | bfela | lgape |
| drame | CCVCV | plare | trale | pluve | trole | rlaep | rteal | vpelu | lrtoe |
| glace | CCVCV | crure | trove | chule | fraga | rruce | vtore | hlecu | rgafa |
| plume | CCVCV | frabe | trifu | vrale | trise | fbate | rftui | eavlr | rstei |
| chien | CCVVC | trouf | crous | treur | drais | rfuot | uosrc | rrteu | rdsia |
| fleur | CCVVC | crior | flain | claut | frout | rcoir | lniaf | ltuac | uoftr |
| fruit | CCVVC | cleur | chuit | cluin | treul | ucrle | htiuc | lciun | lruet |
| train | CCVVC | chaul | stoux | prait | breux | hluac | txuos | rpiat | rxueb |
| jambe | CVCCV | tulbe | torme | madru | vervi | letbu | teomr | adrmu | ervvi |
| larme | CVCCV | butre | fotte | ravre | covre | etbru | tteof | raerv | rvoce |
| sucre | CVCCV | bivre | lebre | sague | mible | rveib | lrbee | gesae | ibmle |
| tigre | CVCCV | vatre | gacle | varte | tiche | ratve | cglea | aervt | cihte |
| vache | CVCCV | sarve | salpe | dutte | techu | evsra | lsape | tdute | hctue |
| villa | CVCCV | nadre | tivre | berde | roste | adrne | rtiev | eebdr | eosrt |
| canal | CVCVC | bulan | zoler | bacal | tomel | bnual | rzeol | cblaa | mloet |
| divan | CVCVC | vadin | ridan | meton | badon | nvdai | dnira | tmeon | dboan |
| roman | CVCVC | nitar | torin | ledin | nimer | iatrn | ntior | ndile | mrnie |
| sapin | CVCVC | barin | palel | fatil | vucor | rnbia | ealpl | lftai | rvocu |
| genou | CVCVV | rabou | tecou | darai | nague | uaorb | euotc | ridaa | gnaeu |
| radio | CVCVV | virie | pomue | bario | tovui | verii | mpeue | rbaio | vtore |
| neige | CVVCV | toudu | bauxe | tiafe | faule | tuudo | xuabe | ftaei | lfuae |
| nuage | CVVCV | naupi | noine | mouse | gouba | npiua | ieonn | smeuo | oagbu |
| piano | CVVCV | vionu | houli | daune | suine | nvoiu | oiulh | dnuea | nseiu |
| arbre | VCCCV | erbra | irdre | onste | inche | eabrr | rrdie | esnto | nhiec |
| oncle | VCCCV | onfle | ancte | onche | ustra | fnloe | acnte | hcnoe | tsrua |
| enfer | VCCVC | erfen | egren | ancal | astin | rfnee | egrne | aanlc | sntai |
| orage | VCVCV | opule | iroge | otire | adite | ueolp | eoirg | rteoi | dtiae |
| usine | VCVCV | orame | ivene | irote | ugone | aomre | vneei | ioetr | uoeng |
| avion | VCVVC | eruit | arous | enait | ocien | rtuei | uraos | itaen | ncioe |

**Figure S1. Individual Z-scores for word-selective responses**

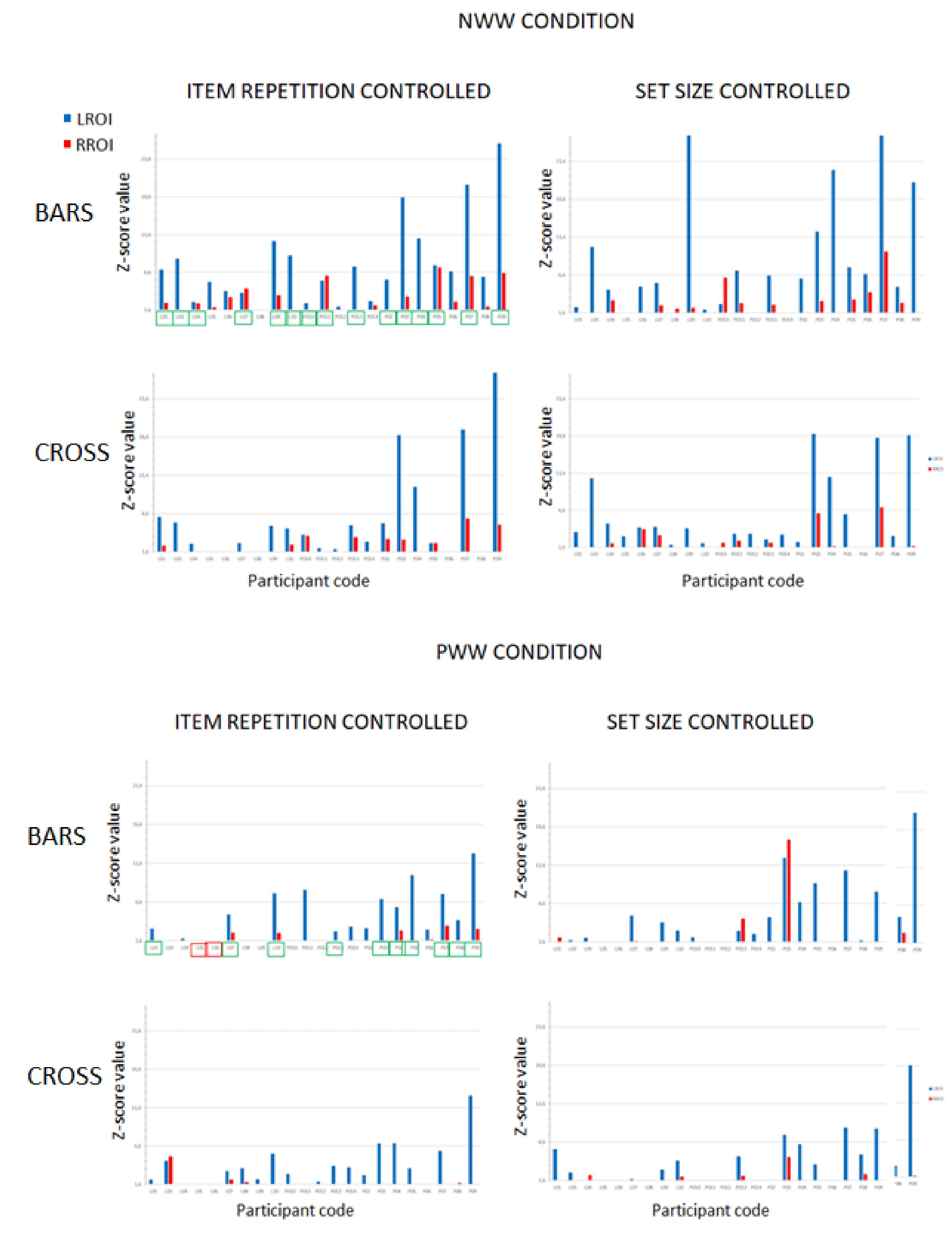


Note: **Z-score values for the words discrimination response per participant**, in the LROI (blue) and RROI (red) in each *discrimination level* (Top: coarse level, words among nonwords NWW; Bottom: fine-grained level, words among pseudowords, PWW), each *Control Type* (item repetition, left columns; set size, right columns) and *Tasks* (bars/ cross). Z-scores were calculated on raw amplitude values after computing the sum of 4 harmonics (2Hz – 8Hz) for the discrimination responses. The lowest Y-axis value has been set to *Z*=1.64 (p<.05, 1-tailed). The first top left histogram for each discrimination level indicates the participants who displayed a significant response *in all* tasks and Control Types (green boxes; 15/22 for NWW and 10/22 for PWW) and in *none* (red boxes; 2/22 for PWW).

**Figure S2. Number of participants with a significant response in the left ROI: A.** Per task x condition (each data point represent 3 minutes testing. In total, there are 76/88 significant data points for NWW, and 58/88 for PWW); **B.** Per task, in *any* condition (at least a significant response in set size *OR* item repetition controlled); **C.** Per task, in *both* conditions (the same individual has a significant response in both set size *AND* item repetition controlled).

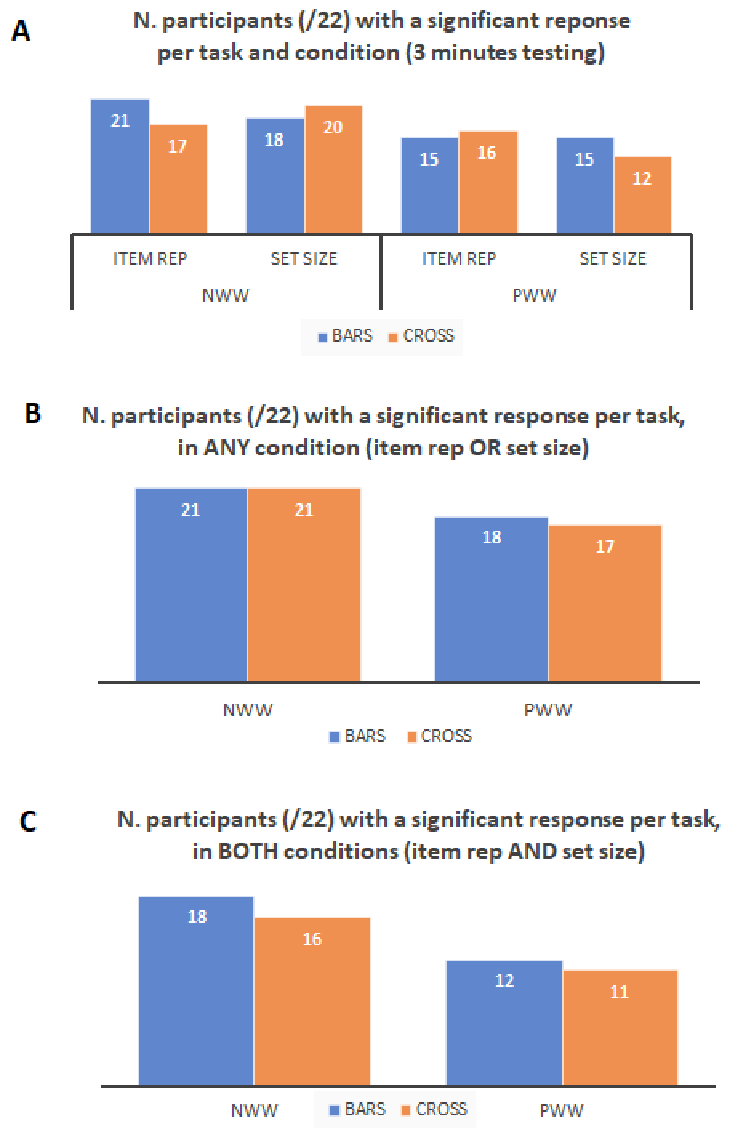
